# NK cells and CTLs are required to clear solid tumor in a novel model of patient-derived-xenograft

**DOI:** 10.1101/2020.05.24.112722

**Authors:** Duy Tri Le, Bryan Burt, George Van Buren, Shawn Abeynaike, Cristina Zalfa, Rana Nikzad, Farrah Kheradmand, Silke Paust

## Abstract

Existing patient-derived-xenograft (PDX) mouse models of solid tumors lack a fully tumor-donor matched “syngeneic” and functional immune system. We developed such a model by engrafting lymphopenic recipient mice with a fresh undisrupted piece of solid tumor, whereby tumor-infiltrating lymphocytes (TIL) expanded in the recipient mice for several weeks. Tumors engrafted in about seventy to eighty percent of <u>s</u>yngeneic-<u>i</u>mmune-<u>s</u>ystem-PDX (SIS-PDX) mice, which harbored tumor-exhausted immune-effector and functional immune-regulatory cells persisting for at least six-months post-engraftment. Interleukin-15 (IL-15)-stimulation in addition to immune checkpoint inhibition (ICI), prevented resistance, resulting in complete or partial response to combined treatment. Further, the depletion of Cytotoxic T lymphocytes (CTLs) and/or Natural Killer (NK) cells from combined immunotherapy in SIS-PDX mice revealed that both cell types are required for the maximal response to tumor. Our novel SIS-PDX model provides a valuable resource for powerful mechanistic and therapeutic studies designed to eradicate solid tumors.

## Introduction

Mounting experimental evidence has documented several key roles for individual and collaborative effector functions of CTLs and NK cells in host immune responses to cancer (*1*–*9*). Activation of CTLs and NK cells have been linked to protective tumor immune surveillance, and higher numbers of tumor-infiltrating CTLs and/or NK cells are favorable prognostic indicators for many types of solid tumors (*10*), including lung cancer (*11*). T cell receptor (TCR) ligation by tumor-derived antigens when presented on the class I human leukocyte antigen (HLA) molecules, in combination with co-stimulatory receptors, activates and expands CTL, which secrete cytokines and chemokines and kill tumor cells (*12*). NK cells express a large number of activating and inhibitory receptors that mostly bind to self MHC-I (*13*, *14*). This is essential in the context of cancer recognition, as MHC-I is often downregulated by malignant cells, circumventing CTL-mediated killing (*15*). Further, tumor-cell-expressed stress ligands can trigger NK cell activating receptors (*14*, *16*, *17*), such as NKG2D (*18*), and induce the secretion of cytokines and chemokines, as well as pore-forming proteins (perforin) and cytotoxic mediators (granzymes) that trigger tumor-cell apoptosis (*19*).

Immune suppressive cells, *e.g.,* myeloid derived suppressor cells (MDSC) (*20*), M2 macrophages (*21*, *22*), and regulatory T cells (Tregs) (*23*), inhibit CTL- and NK cell-effector function in solid tumors (*24*) via the expression of inhibitory ligands, suppressive cytokines, and tumor-promoting factors (*25*). Indeed, the abundance of MDSC and Tregs in solid tumors positively correlate with advanced disease and increased tumor burden (*26*–*28*), and are independent predictors of poor outcome (*29*–*32*). Concurrently, pharmacological targeting of MDSC and Tregs in animal models and clinics significantly improves anti-tumor immunity enabling tumor control (*33*, *34*). In addition to the suppression of cytotoxic anti-tumor activities by immune suppressive cells, tumors further trigger cell-intrinsic defects and the upregulation of immune checkpoint molecules in CTLs and NK cells via chronic overstimulation (*35*).

Immune checkpoint molecules have several non-redundant roles and distinct expression patterns that are required to prevent autoimmunity under normal conditions. In support of this concept, immune-checkpoint-deficient mice display auto-inflammatory diseases in multiple organs (*36*). A well-recognized mechanism for tumor evasion is the induction of immune checkpoint molecules such as programmed death (PD) and its ligands (PD-L) that prevent efficient T and NK cell activation (*37*). Indeed, tumor-exhausted CTLs and NK cells express high levels of PD-1 (*37*), and transformed cells escape immune attack by expressing significant levels of PD-ligands (*38*). ICI targeting of immune suppressive pathways are FDA approved for the treatments of many solid tumors, including non-small-cell-lung cancer (NSCLC). However, blocking immune-checkpoint molecules alone does not guarantee tumor eradication by cytotoxic immune cells, as immune cell attack ultimately depends on the recognition of either tumor antigen by T cells, or stress-ligand-mediated activation of NK cells (*39*–*41*). This may explain why ICI is most effective in tumors with a high metastatic burden, which presumably results in an abundance of tumor antigens that are stimulatory to the immune system (*42*). Lung cancer is responsible for more than one quarter of all cancer deaths worldwide (*43*). Lung adenocarcinomas (LUAD), a subtype of NSCLC, is a genetically heterogeneous cancer that is in part the result of oncogenic mutations and the loss of tumor suppressors (*44*–*48*). The Food and Drug Administration (FDA) has approved PD-1 or PD-L1 blockade for advanced NSCLC treatment (*24*, *31*, *48*, *49*), albeit it is not a cure (*50*–*52*).

Existing PDX models lack a human tumor-donor-matched, complete, functional and mouse-tolerant immune system (*53*–*57*) and, consequently, lack the utility for specific types of cancer immunotherapy and mechanistic studies. To fulfill this unmet need, we developed a PDX model of human solid tumor with a syngeneic complete mouse tolerant and functional immune system that reconstitutes recipient mice for at least six months post tumor engraftment, is responsive to combination immunotherapy and allows for in depth mechanistic *in vivo* studies. Specifically, using our novel PDX model, we evaluated the efficacy of combination immunotherapy on the eradication of lung cancer and identified immune cells required for therapy success. Our novel PDX model delivers a valuable resource for powerful pre-clinical mechanistic and therapeutic studies to eradicate solid tumor.

## Results

### NK cells in LUAD display exhausted markers

We first used flow cytometry to identify the NK cell phenotypes of tumor-infiltrating and donor-matched uninvolved lung-tissue and PBMC-derived NK cells. We found that NK cell frequencies were significantly decreased in LUAD, and fewer LUAD-resident NK cells expressed the activating receptors CD16 and/or NKG2D compared to donor matched lung tissue- or PBMC-derived NK cells (**Figure 1A, B, D, E**). Notably, tumor-resident NK cells expressed significantly higher PD-1 and/or both of its’ ligands, PD-L1 and PD-L2 when compared to syngeneic lung- or PBMC-derived NK cells (**Figure 1C**), while PB-derived NK cells did not express PD-1, PD-L1 or PD-L2 (**Figure 1 F**). We unexpectedly found high numbers of PD-L1 and PD-L2-expressing NK cells in both non-tumor lung-tissue and lung-tumor, however, this was consistent across genetically unrelated human donors (**Figure 1C**). These data suggest that PD-1, PD-L1 and PD-L2 are expressed on lung tissue- as well as lung-tumor-resident NK cells, and a significantly higher number of NK cells express PD, PD-L1 and/or PD-L2 in tumor compared to lung-tissue.

**Figure 1.**
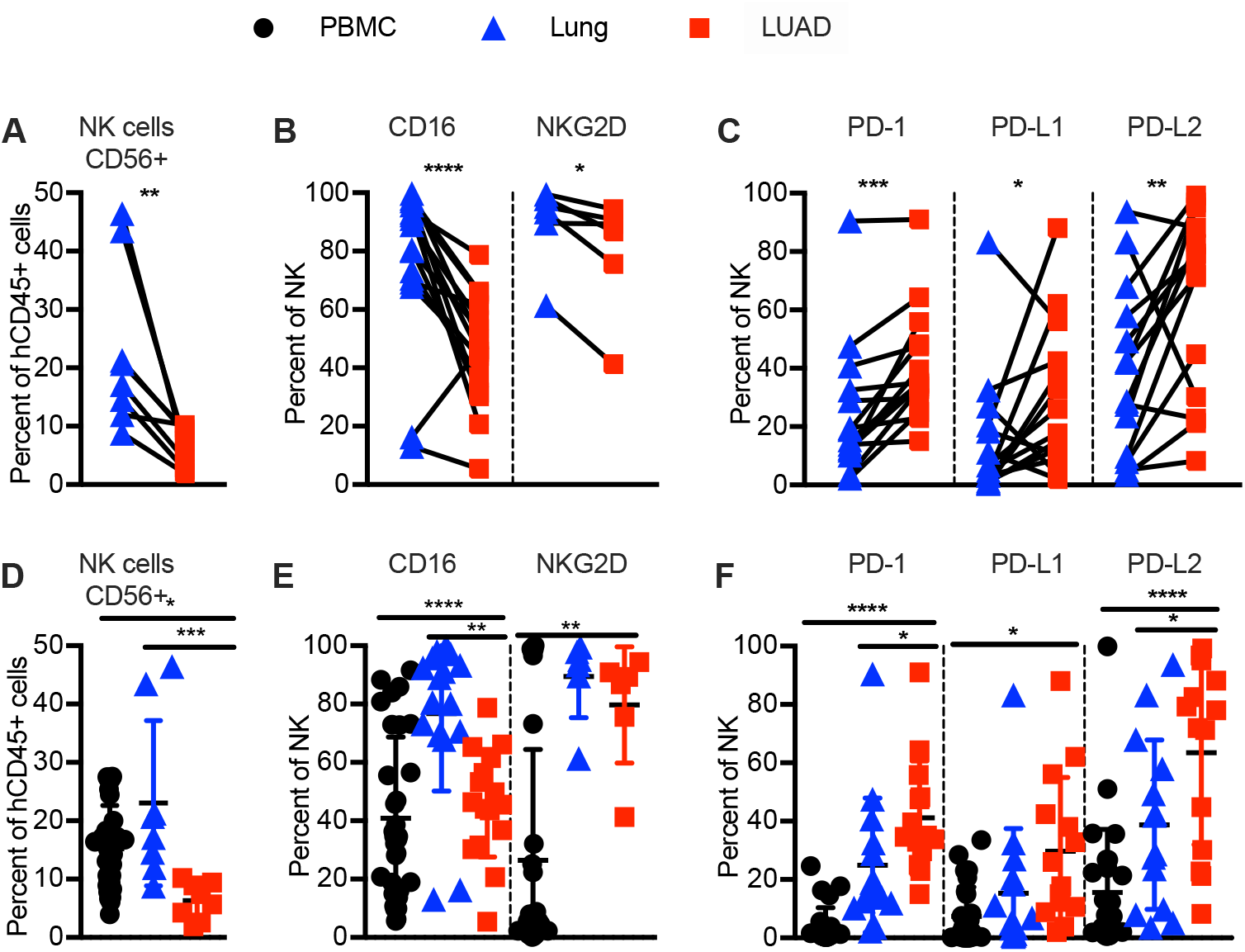
Human LUAD associated NK cells are exhausted in phenotype. Immune phenotyping of human NK cells in indicated tissues by flow cytometry. (**A, D**) Frequency of human NK cells (CD45^+^CD3^−^CD56^+^) in donor-matched lung vs. NSCLC (LUAD) tissue. (**B, E**) Frequency of CD16 and/or NKG2D expressing human NK cells, (**C, F**) Frequency of PD-1, PDL-1 and/or PDL-2-expressing human NK cells in donor-matched lung-tissue vs. LUAD (**A-C, top**), or LUAD-patient derived PBMC, lung tissue and LUAD (**D-F, bottom**). Each data point represents one genetically unrelated human donor (N= eight to 31 depending on tissue). Paired t-test for donor-matched lung tissue vs. LUAD, one-way ANOVA with post-hoc Tukey test for comparisons between LUAD-patient derived PBMC, lung tissue and LUAD (not always donor matched between all three tissues). *p < 0.05; **p < 0.01; ***p < 0.001, ****p<0.0001

### T cells in LUAD display exhausted markers

While the numbers of hematopoietic cells in lung tissue and LUAD were similar (**Figure 2A**), T cells, in contrast to NK cells, were more frequent in tumor-tissue vs. syngeneic lung-tissue (**Figure 2 B, D, F, G, I**). However, this was due to an increase in the immune suppressive tumor-resident Treg subset (**Figure 2 E, J**). As expected, PD-1 expression was increased in lung-tumor-infiltrating T cells compared to donor matched lung-tissue, and T cells in both lung-tissue and tumors expressed significantly higher levels of PD-L2 (**Figure 2 C**). PBMC-derived T cells did not express significant levels of PD family members, including either PD ligand (**Figure 2 H**), suggesting that T cells, similar to NK cells, express PD-1 and/or PD-ligands when resident in lung tissue and/or lung tumor, but not when in PBMC.

**Figure 2.**
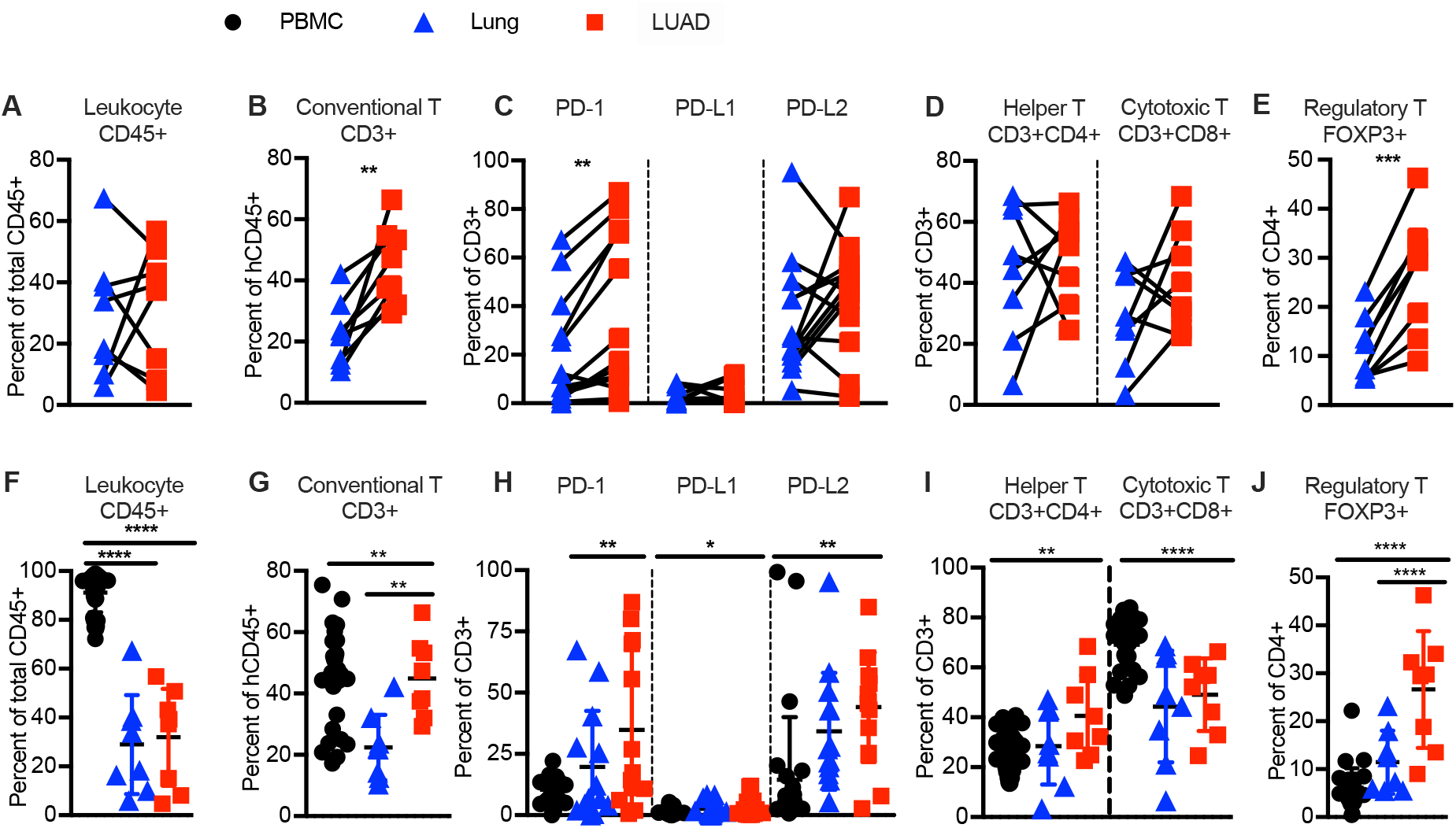
Human T cells are exhausted in phenotype and enriched in T regulatory cells in NSCLC (LUAD). Immune phenotyping of human immune cells and T cell subsets in indicated tissues by flow cytometry. (**A, F**) Frequencies of human hematopoietic cells (human CD45^+^), (**B, G**) conventional T cells (CD45^+^CD3^+^CD56^−^), (**C, H**) PD-1, PD-L1 and/or PD-L2 expressing conventional T cells, (**D, I**) T helper (CD45^+^CD3^+^CD4^+^CD8^−^CD56^−^), and CTL CD45^+^CD3^+^CD4^−^ CD8^+^CD56^−^subsets, (**E, J**) T-regulatory cell (CD45^+^CD3^+^CD4^+^Foxp-3^+^CD8^−^CD56^−^) subset in donor-matched lung vs. NSCLC (LUAD) (**A-E, top**), or patient derived PBMC, lung tissue and LUAD (**F-J, bottom**). Each data point represents one genetically unrelated human donor (N= 8-31 depending on tissue, same donors as for Figure 1). Paired t-test for donor-matched lung tissue vs. LUAD, one-way ANOVA with post-hoc Tukey test for comparisons between LUAD-patient derived PBMC, lung tissue and LUAD (not always donor matched between all three tissues). *p < 0.05; **p < 0.01; ***p < 0.001, ****p<0.0001

### Generation of donor-matched LUAD-SIS-PDX-immune system mice

To generate our novel syngeneic model of SIS-PDX, we transplanted an un-disrupted piece of freshly resected LUAD from newly diagnosed and treatment-naïve subjects into a skin pocket of each recipient NSG mouse (**Figure 3A**). About six weeks later, LUAD had engrafted in about 70% of NSG mice with extensive vascularization, when human donors and recipient NSG mice were sex matched. Histological analyses revealed carcinoma architecture with hyperproliferating foci (**Figure 3B**). We did not transfer any patient derived PBMC or stem cells, or any cells from sources other than the tumor, but instead tested the breath and longevity of TIL-derived immune cell reconstitution in LUAD-SIS-PDX mice. We used flow cytometry to determine the frequencies of human hematopoietic cells, NK cells, conventional T cells, including T helper, CTL, and Treg, one-, two- and six-months post tumor transplant. We found that all of these immune cell types were present in PDX mice for at least six months (**Figure 3 C-E**). As in humans, NK cells made up 5-10% of peripheral blood (PB) hematopoietic cells in LUAD-SIS-PDX mice, with a predominance of human T lymphocytes. T cells were composed of CD4^+^ T helper and CD8^+^ T CTL at a 2:1 ratio. Additionally, B cells and dendritic cells were also present in LUAD-SIS-PDX mice (**Supplemental Figure 1**), demonstrating long-term reconstitution with full syngeneic human immune cells and underscoring the successful development of a PDX model in which immune cells are donor matched to the tumor. Notably, most LUAD-SIS-PDX from a single donor developed robust reconstitution with a complete TIL-derived immune system in a cohort of 12-15 mice (**Figure 4**). As tumors generally become palpable and start growing noticeably around four to six weeks post-transplant, this immune cell reconstitution timeline allows for several months of immunotherapy studies in our novel LUAD-SIS-PDX model.

**Figure 3.**
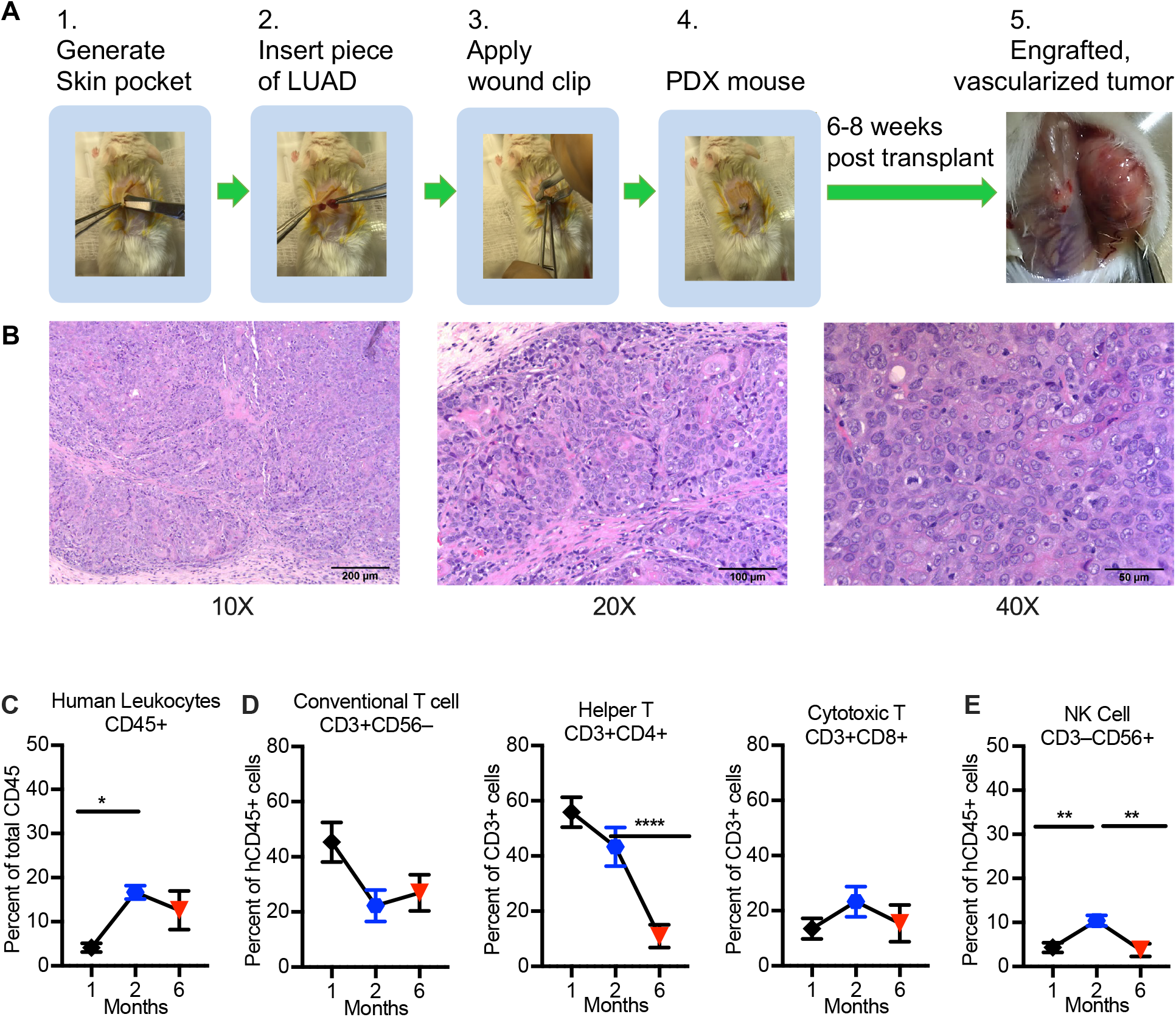
Human immune cells persist for months in PDX mice. (**A**) Generation of PDX mice by implantation of human LUAD into a skin pocked of NSG recipient mice. (**B**) Representative H & E-staining of PDX-derived LUAD 14 weeks post-transplant demonstrating retained tumor architecture (**C-E**) Flow cytometry of human immune cells from PBMC of PDX mice, 1-month, 2-months, or 6-months transplant. Percentage of (**C**) human hematopoietic cells (human CD45^+^) of total (mouse + human) hematopoietic cells, (**D**) conventional T cells (CD45^+^CD3^+^CD56^−^), including T helper (CD45^+^CD3^+^CD4^+^CD8^−^CD56^−^), and CTL CD45^+^CD3^+^CD4^−^CD8^+^CD56^−^ subsets, and T regulatory cells (CD45^+^CD3^+^CD4^+^Foxp-3^+^CD8^−^CD56^−^), as well as (**E**) NK cells (CD45^+^CD3^−^CD56^+^) at indicted times post-transplant (N = 15-23 PDX mice per group from six genetically unrelated donor-cohorts). Data are represented as mean SEM; one-way ANOVA with post-hoc Tukey test *p < 0.05; **p < 0.01; ***p < 0.001.

**Figure 4.**
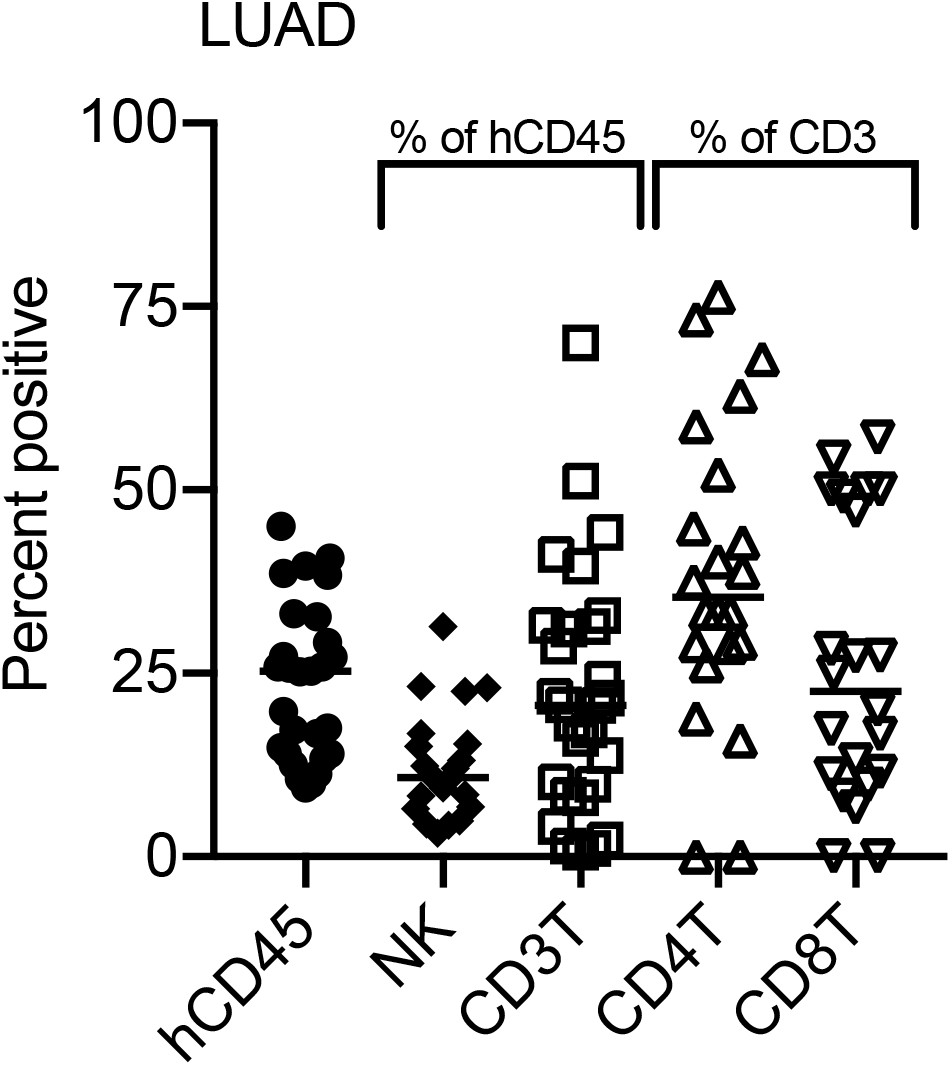
Immune cell reconstitution across a cohort of LUAD-SIS-PDX mice 4 months post engraftment. Flow cytometry of human immune cells from PBMC of single LUAD-PDX cohort of 15 mice analyzed 4-months post-transplant. Frequency of human hematopoietic cells (human CD45^+^ of total CD45^+^), NK cells (CD45^+^CD56^+^CD3^−^), NKT cells (CD45^+^CD56^+^ CD3^+^), conventional T cells (CD45^+^CD56^−^CD3^+^), T helper cells (CD45^+^CD56^−^CD3^+^CD4^+^ CD8^−^), and Cytotoxic T cells (CD45^+^CD56^−^CD3^+^CD8^+^CD4^−^). Each data point represents one LUAD-SIS-PDX mouse of a single human donor cohort, N=15 animals.

### Human NK and T cells in LUAD-SIS-PDX mice resemble exhausted TILs of LUAD donors

To evaluate whether immune exhaustion to tumor is preserved on NK and T cells of LUAD-SIS-PDX mice, we evaluated the immune phenotypes of human NK and T cells in mouse PB, spleens and engrafted human tumor using multiparametric flow cytometry (**Figure 5**). Hematopoietic cells (**Figure 5 A**) and NK cells (**Figure 5 B**) were present in PB, spleens and tumor of LUAD-SIS-PDX mice, and the expression of the activating receptors CD16 and NKG2D did not significantly differ between tissues (**Figure 5 C**). Further, similar to NK cells from freshly-dissected LUAD-tissue (**Figure 1 D, F)**, more NK cells expressed PD-L2 than PD-L1 (**Figure 5 D**). However, the frequency of PD-L1-expressing NK cells was reduced in PDX-LUAD compared to freshly excised LUAD. Reduced PD-L1 expression was also observed in transplanted non-hematopoietic cells of the tumor compared to freshly excised LUAD, while PD-L2 expression was preserved (**Supplemental Figure 2**). In contrast, human T cells that reconstitute peripheral organs of LUAD-SIS-PDX mice resembled T cells in freshly excised LUAD tissue (**Figure 2 C, H**) in their expression of PD-1 and PD-L2, with a lack of PD-L1 expression by T cells (**Figure 5 E-H**). As our freshly excised and SIS-PDX derived tumor tissues are not always from the same donor, it is not possible to determine whether this reduction in PD-L1 expression is due to patient to patient variation or whether factors that drive PD-L1 expression on human NK cells and LUAD in humans are reduced in PDX mice. We conclude that LUAD-resident NK and T cells survive long-term in PDX mice, in which they repopulate peripheral tissues and largely maintain their expression of markers of exhaustion.

**Figure 5.**
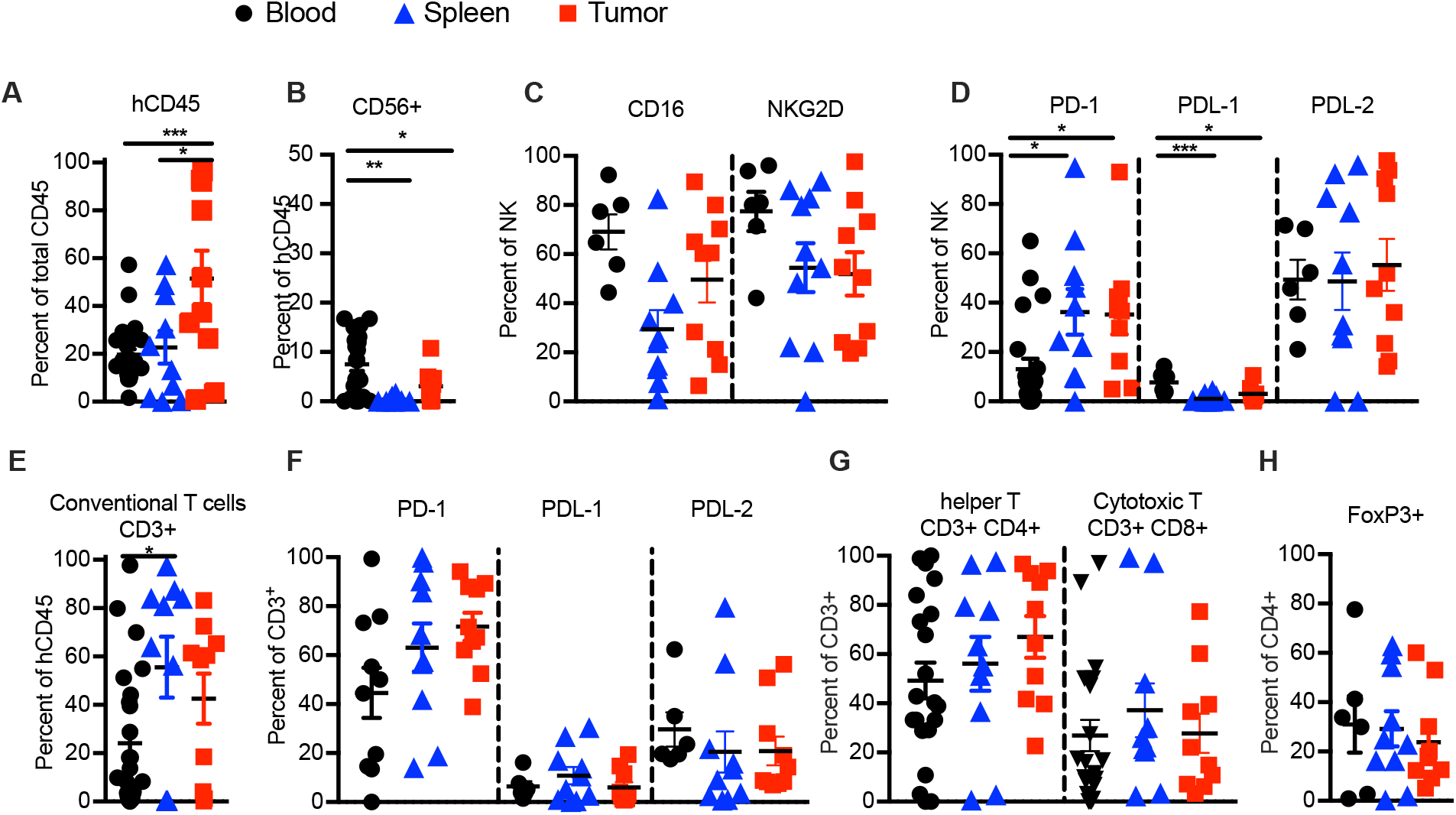
Immune cells in LUAD-SIS-PDX mice resemble exhausted Tumor Infiltrating Immune cells (TILs) cells of LUAD donors. (**A-H**) Flow cytometry of human immune cells from PBMC, spleen, and transplanted LUAD of PDX mice (“tumor”) analyzed 2-months post-transplant. (**A**) Frequency of human hematopoietic cells (CD45^+^), (**B**) NK cells (CD45^+^CD3^−^ CD56^+^), and (**C**) expression of activating receptors CD16 and NKG2D on NK cells, and (**D**) PD-1, PD-L1 and PD-L2 on NK cells in PDX mouse derived PBMC, spleen and tumor. (**E**) Frequency of conventional T cells (CD45^+^CD3^+^CD56^−^), and (**F**) expression of PD-1, PD-L1 and PD-L2 on conventional T cells, as well as (**G**) frequencies of T helper (CD45^+^CD3^+^CD4^+^CD8^−^ CD56^−^) and CTL CD45^+^CD3^+^CD4^−^CD8^+^CD56^−^), and (**H**) T-reg (CD45^+^ CD3^+^CD4^+^Foxp-3^+^CD8^−^ CD56^−^) subsets in PDX mouse derived PBMC, spleen and tumor. (N = 10 - 23 PDX mice per group from five genetically unrelated human donor cohorts). Data are represented as mean SEM; one-way ANOVA with post-hoc Tukey test *p < 0.05; **p < 0.01; ***p < 0.001.

### Establishment of the SIS-PDX model with pancreatic adenocarcinoma cancer (PDAC)

PDAC is high in immune suppressive T-reg and MDSC (*58*), and it’s fibrous stroma expresses chemokines that trap CTL (*59*–*61*). Due to its rather intermediate tumor mutation burden, PDAC is poorly immunogenic (*42*) and has shown little susceptibility to ICI (*62*, *63*), and is the 4^th^ leading cause of cancer death in western countries (*64*). To support the future development of improved immunotherapies for PDAC, we evaluated whether PDAC is suitable for the generation of SIS-PDX mice. To do so, we engrafted NSG mice each with a piece of treatment-naïve pancreatic duct adenocarcinoma (PDAC) tissue (**Figure 6**), and evaluated subsequent tumor growth, immune cell reconstitution and expression of checkpoint molecules on T and NK cells by flow cytometry, four to six months post tumor engraftment. Immune phenotyping of NK cells in freshly excised pancreas and PDAC, and comparisons to PBMC revealed that NK significantly fewer NK cells express the activating receptor CD16 in pancreas and PDAC compared to PBMC. Further, while NKG2D-expressing NK cells were abundant in pancreas, they were rare in PBMC and PDAC (**Figure 6C**). Similar to PD-1 and PD-L2 expression on lung tissue resident NK cells, pancreas tissue resident NK cells expressed PD-1 and PD-L2. In contrast to lung tissue however, PD-L1 expression was not different between PBMC, pancreas and PDAC -resident NK cells (**Figure 6D**). Interestingly, T cells expressed PD-1 but not PD-ligands in all three tissues, and, similar to LUAD, were increased in frequency in tumor due to a significant increase in Treg (**Figure 6 E-H**).

**Figure 6.**
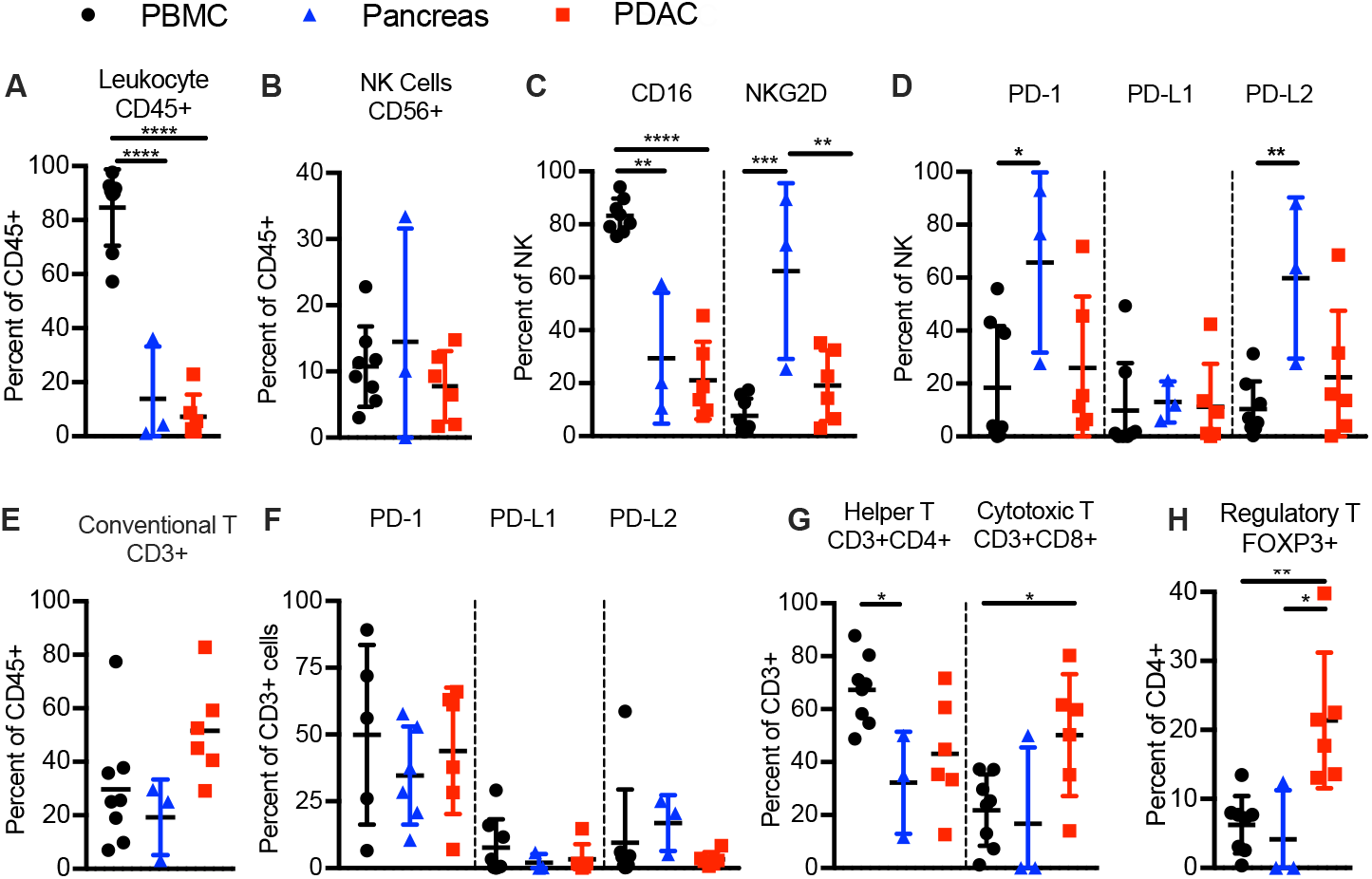
Human PDAC associated NK and T cells are exhausted in phenotype. Frequency of (**A**) human hematopoietic cells (human CD45^+^), (**B**) NK cells (CD45^+^CD56^+^CD3^−^), (**C**) CD16 and/or NKG2D expressing human NK cells, (**D**) PD-1, PDL-1 and/or PDL-2-expressing human NK cells in in indicated freshly excised tissues of PDAC patients. Frequency of (**E**) human conventional T cells (CD45^+^CD56^−^CD3^+^), (**F**) PD-1, PDL-1 and/or PDL-2-expressing human T cells in PDAC-patient derived PBMC, pancreas and PDAC. Frequency of (**G**) Th (CD45^+^CD3^+^CD4^+^ CD8^−^) and CTL (CD45^+^CD3^+^CD4^−^CD8^+^) as percent of human T cells, and (**H**) Treg (CD45^+^CD3^+^CD4^−^CD8^+^Foxp3^+^) as percent of Th cells in PDAC-patient derived PBMC, pancreas and PDAC. Each data point represents one genetically unrelated human donor. Paired t-test for donor-matched lung tissue vs. PDAC, one-way ANOVA with post-hoc Tukey test for comparisons between PDAC-patient derived PBMC, pancreas tissue and PDAC. *p < 0.05; **p < 0.01; ***p < 0.001, ****p<0.0001

Similar to LUAD-SIS-PDX mice, PDAC-SIS-PDX mice harbored human hematopoietic cells (**Figure 7A**), including effector cells such as NK cells (**Figure 7 B-D**), and conventional T cells (**Figure 7 E-H**), including Th, CTL (**Figure 7 G**) and Tregs (**Figure 7H**). In addition to effector cells and Treg, IL-10 and/or TGF-beta-producing MDSC (**Supplemental Figure 3 A-C**) and M2 monocytes (**Supplemental Figure 3 D-F**) were present in PDAC-SIS-PDX PB and peripheral tissues four- to six- months post tumor engraftment. PDAC-SIS-PDX-resident NK cells expressed varying levels of CD16, NKG2D, PD-1, PD-L1 and PD-L2, and their expansion in PDAC-SIS-PDX mice did not rescue their exhausted phenotype (**Figure 7 A-D**). Similarly, T cells, including significant amounts of T-reg, expanded in PDAC-SIS-PDX mice, and expressed PD-1, but not PD-L1 or PD-L2, resembling PDAC-donor tissues (**Figures 7 E-G**). Also preserved were PD-L1 and PD-L2 expression on PDAC, as freshly excised PDAC expressed PD-L1 and PD-L2, and expression was maintained in PDAC-SIS-PDX (**Supplemental Figure 4**). We conclude that SIS-PDX mice generated with LUAD or PDAC reconstitute a tumor-exhausted immune system resembling that of the human donor, and that this immune reconstitution lasts for several months.

**Figure 7.**
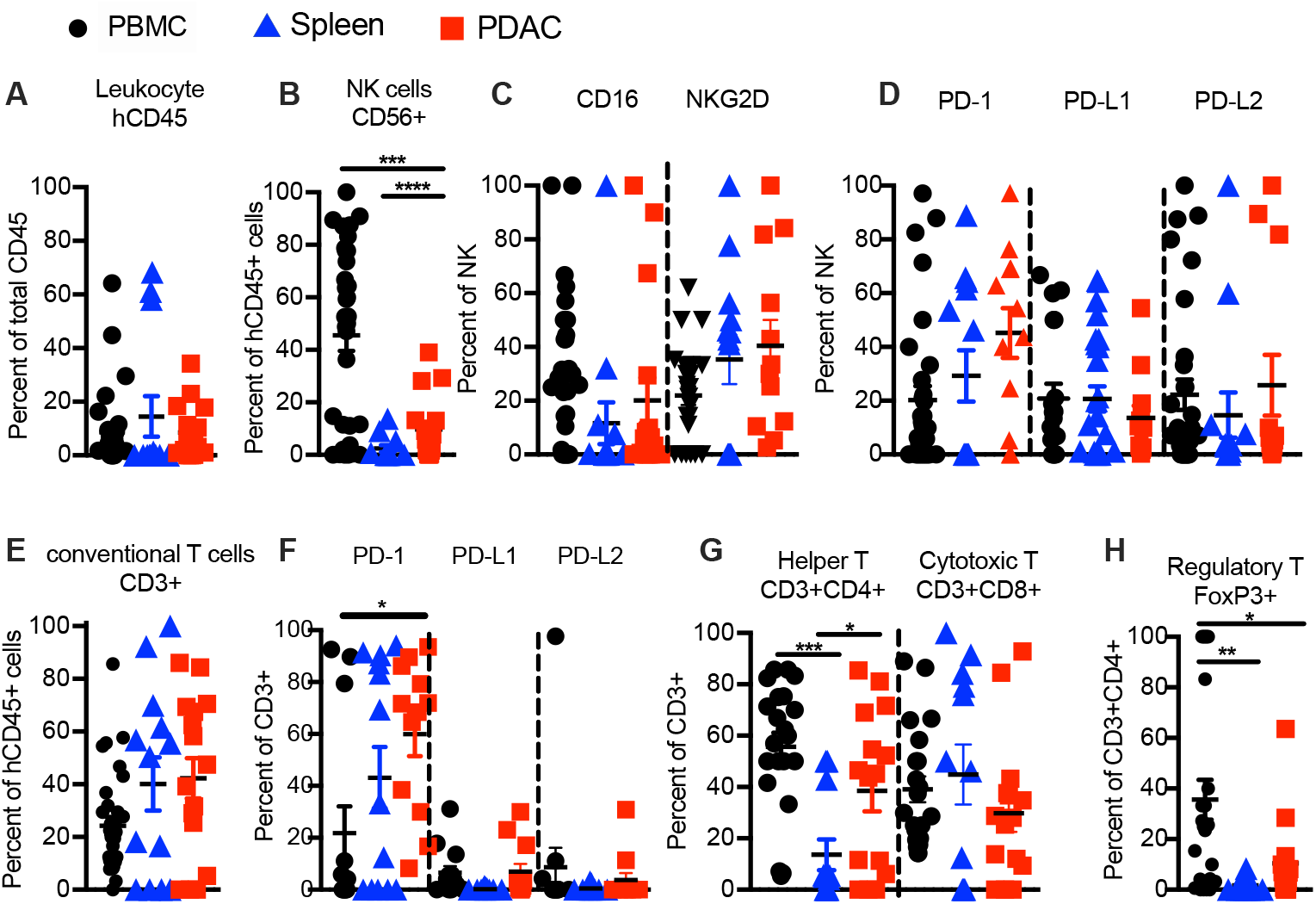
Immune cells in PDAC-SIS-PDX mice resemble exhausted TILs of PDAC donors. (**A-H**) Flow cytometry of human immune cells from PBMC, spleen, and transplanted PDAC of PDX mice analyzed 4-6 months post-transplant. (**A**) Frequency of human hematopoietic cells, (**B**) human NK cells (CD45^+^CD56^+^CD3^−^), and (**C**) expression of activating receptors CD16 and NKG2D as well as (**D**) markers of exhaustion PD-1, PD-L1 and PD-L2 on NK cells in PDX derived PBMC, spleen and transplanted PDAC. (**E**) Frequency of conventional T cells (CD45^+^CD56^−^CD3^+^) and (**E**) expression of PD-1, PD-L1 and PD-L2 on conventional T cells, (**G**) frequencies of Th (CD45^+^CD3^+^CD4^+^CD8^−^), and CTL (CD45^+^CD3^+^CD4^−^CD8^+^) as percent of T cells, and (**H**) Treg (CD45^+^CD3^+^CD4^−^CD8^+^Foxp3^+^) as percent of Th cell subsets in PDX derived PBMC, spleen, and transplanted PDAC. Each data point represents one PDAC-SIS-PDX mouse tissue (n= 12-33 PDX mice per group, seven donor-cohorts). Data are represented as mean SEM; Paired t-test for donor-matched lung tissue vs. PDAC, one-way ANOVA with post-hoc Tukey test *p < 0.05; **p < 0.01; ***p < 0.001.

### Correlation between tumor growth and immune cells in SIS-PDX model

To determine residual anti-tumor immune function in our novel SIS-PDX model, we next examined the correlation between tumor growth and the presence of tumor-fighting effector cells, and/or immune exhaustion (**Figure 8 A-D**). We found a significant inverse correlation between the presence of human immune cells (**Figure 8 A**), NK cells (**Figure 8 B**) or T cells (**Figure 8 C**) and tumor size, and a significant correlation between NK cell-expressed PDL-1 and PDL-2 and tumor size, while NK cell-expressed PD-1 did not correlate significantly with tumor growth (**Figure 8 D**). These correlations supported an increase T and NK cell infiltration in transplanted tumors with smaller tumors, while a higher expression of NK cell-expressed checkpoint molecules is found in larger tumors. These findings further suggest that CTL and NK cells are active tumor fighters, and that NK cells are susceptible to checkpoint inhibition in LUAD-SIS-PDX mice.

**Figure 8:**
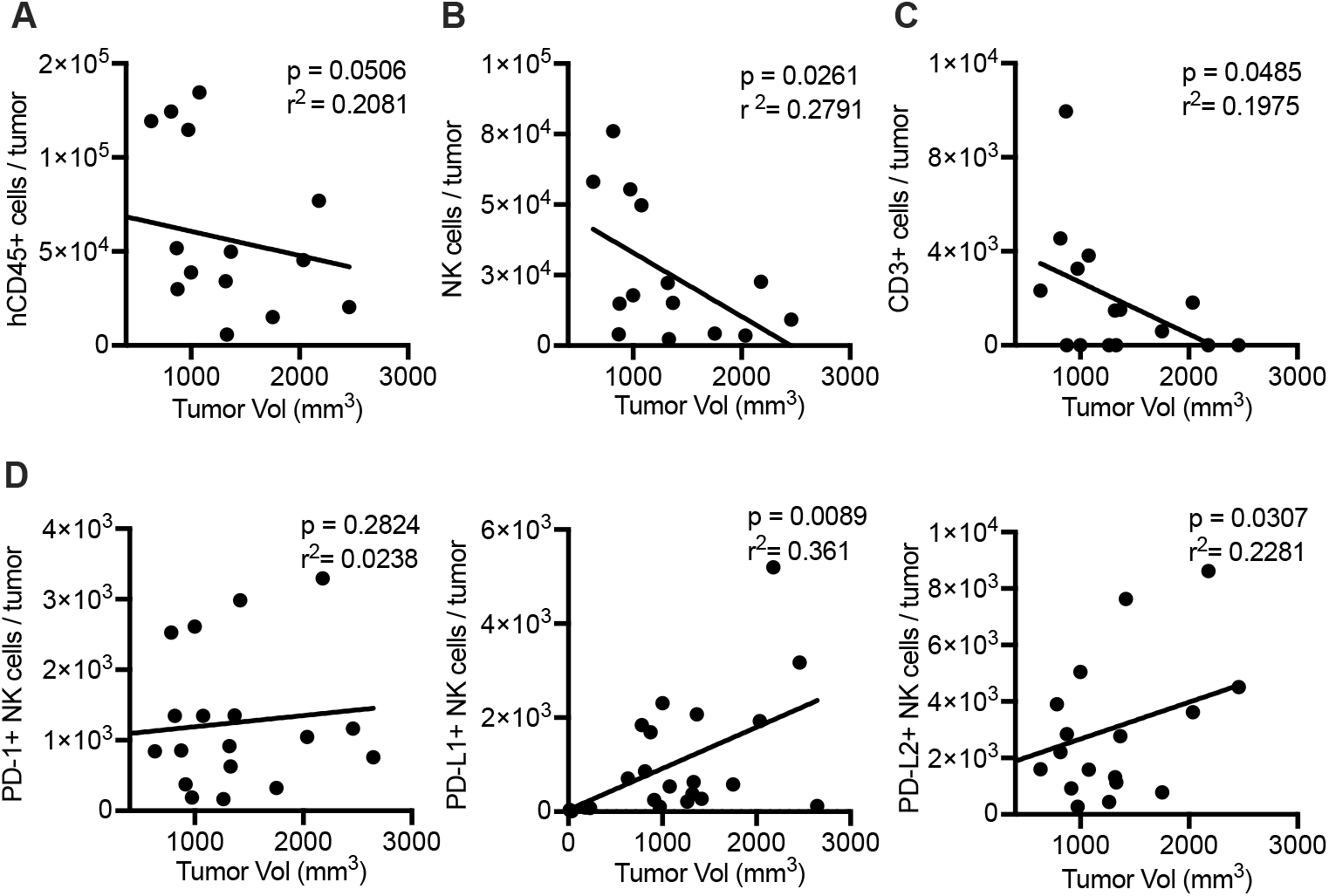
Correlation of numbers of tumor resident human immune cells and tumor size in LUAD-SIS-PDX mice 6 months post-transplant. (**A-D**) Linear regression plots demonstrating correlations between numbers of human (**A**) hematopoietic cells (CD45^+^), (**B**) NK cells (CD45^+^CD3^−^CD56^+^), and (**C**) conventional T cells (CD45^+^CD3^+^CD56^−^), or (**D**) indicated immune-modulatory receptors expressed on NK cells and tumor volumes 6 months post-transplant. N = 14-16 LUAD-SIS-PDX mice were analyzed for each correlation from three genetically unrelated donor cohorts. Correlations were computed using Pearson correlation coefficients, p<0.05 was considered a statistically significant difference.

### IL-15-stimulation prevents tumor escape from ICI resulting in complete or partial remission

We hypothesized that immunotherapy restoring the activation of tumor-exhausted T and NK cells will elicit potent anti-tumor responses in one or both cytotoxic immune cell types. To achieve this goal, we chose a combination of PD-1 blockade and IL-15 stimulation as a combination immunotherapy, as both treatments are reported to be safe for humans (*49*). IL-15-stimulation increases the expression of the activating receptors CD16 and NKG2D on NK cells, in addition to increasing the activation, proliferation, cytotoxic activity and survival of NK cells and CTLs (*65*). We found that the combination of PD-1 blockade and IL-15 signaling resulted in the eradication of transplanted LUAD in about one half of treated LUAD-SIS-PDX mice, while the other half presented with a partial response (**Figure 9**). Notably, trans-presented IL-15 alone, but not PD-1 blockade significantly reduced tumor burden in all treated LUAD-SIS-PDX animals. PD-1 blockade alone transiently prevented tumor growth, but after two weeks, tumors grew at a similar rate to untreated control tumors **(Figure 9)**. The addition of IL-15 to PD-1 blockade completely abrogated tumor escape from ICI, resulting in a powerful additive therapeutic effect capable of tumor eradication. These findings support a key role for adjuvant IL-15 treatment to induce immune cell-mediated tumor attack, which can prevent tumor escape from checkpoint blockade therapy as shown using our novel LUAD-SIS-PDX model.

**Figure 9.**
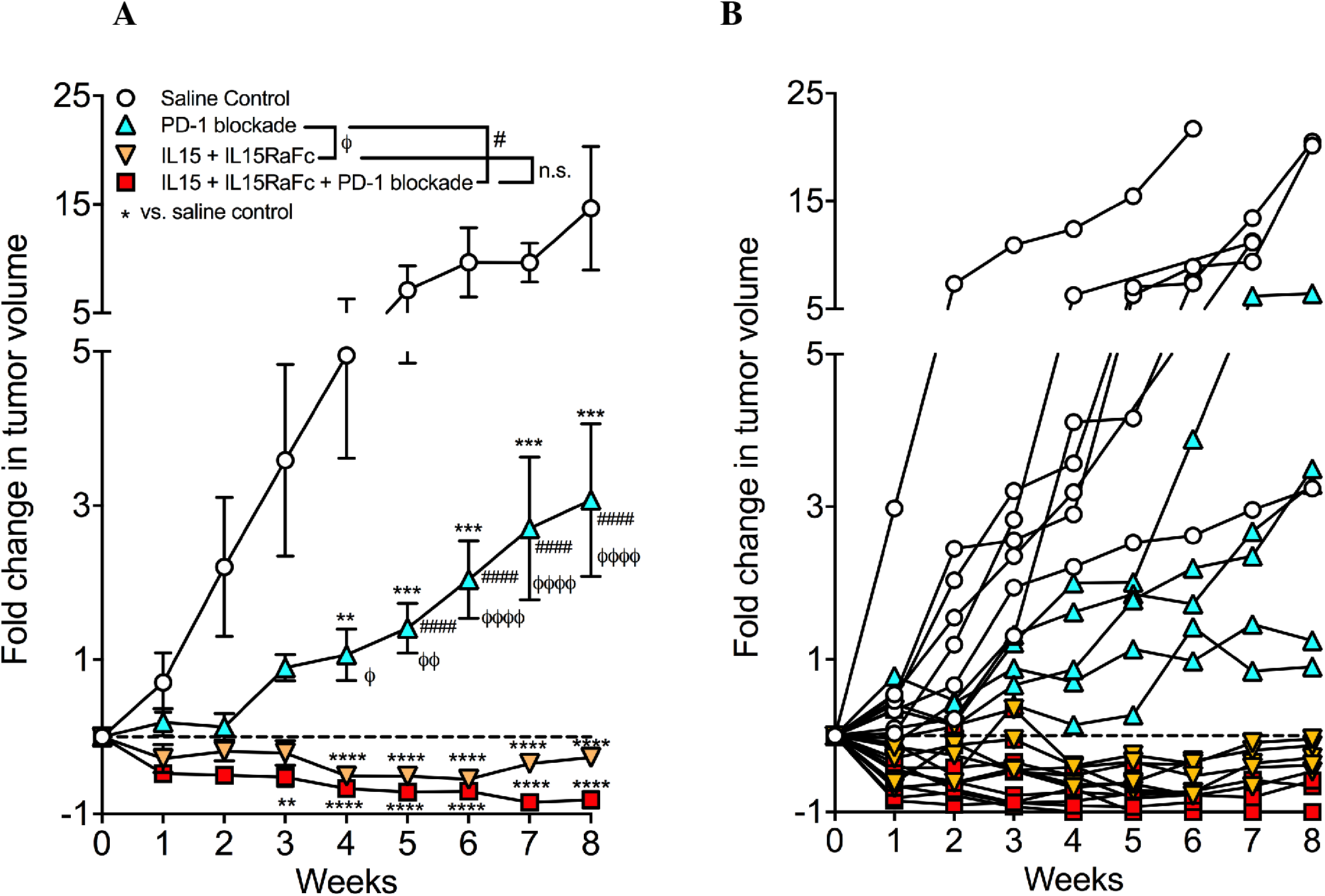
IL-15-stimulation prevents tumor escape from ICI resulting in complete or partial remission. Fold change in tumor volume in LUAD-SIS-PDX mice treated with 100μg/mouse blocking antibody to PD-1, 5μg/mouse IL-15 + 2.5μg/mouse IL-15RaFc, or both PD-1 and IL-15/IL-15RaFc as indicated by intraperitoneal injection at day 0 and every seven days thereafter. Control mice received saline. Experiments were started when tumors were about 50 +/− 14 mm^3^ with no statistic size difference between experimental and control groups at day 0 and tumor volumes measured weekly. (**A**) Mean fold change in tumor volume of each experimental or control group with standard error of the mean. (**B**) Mean fold change in tumor volume of LUAD-SIS-PDX mouse with standard error of the mean. N = 5-11 LUAD-SIS-PDX mice per group from three genetically unrelated human donor cohorts. Data are represented as mean and SEM; Multiple t-test, *,^#^,^ϕ^ p < 0.05; **,^##^,^ϕϕ^ p < 0.01; ***,^###^,^ϕϕϕ^ p < 0.001.

### NK cells and CTLs are required for successful combination immunotherapy in LUAD-SIS-PDX model

We hypothesized that CTLs and NK cells are key targets in combined immunotherapy response in LUAD-SIS- PDX model. To dissect their individual contributions to tumor eradiation, we depleted either CTL, circulating NK or both effector cell types using antibody specific to CD8a (CTL-depletion) or NKp46 (NK cell depletion), and treated LUAD-SIS-PDX mice with IL-15/IL-15RaFc + PD-1-blockade combination immunotherapy. We found that the depletion of either CTL or NK cells resulted in stable disease, while depletion of both cell types unleashed rapid and lethal tumor growth even in the presence of immunotherapy (**Figure 10**). Similarly, when previously frozen tumors were used for implantation, whereby freezing of tumor tissue resulted in the death of TILs, the treatment of tumor-bearing PDX mice with IL-15/IL-15RaFc + PD-1-blockade was ineffective and tumor growth accelerated five-fold (**Supplemental Figure 5**). We conclude that IL-15/IL-15RaFc + PD-1-blockade activates both CTL and NK cells, that CTL and NK cells are the sole targets of this therapy, and that CTL and NK cells are equally required for a maximal anti-tumor response to cure human solid tumor in our syngeneic PDX model of lung cancer. We further conclude that our novel LUAD-SIS-PDX model not only enables much needed efficacy evaluations of immunotherapies to effectively reverse tumor-induced immune suppression, but further permits powerful mechanistic studies that identify immune effector cells crucial to the eradication of solid tumor upon successful immunotherapy.

**Figure 10.**
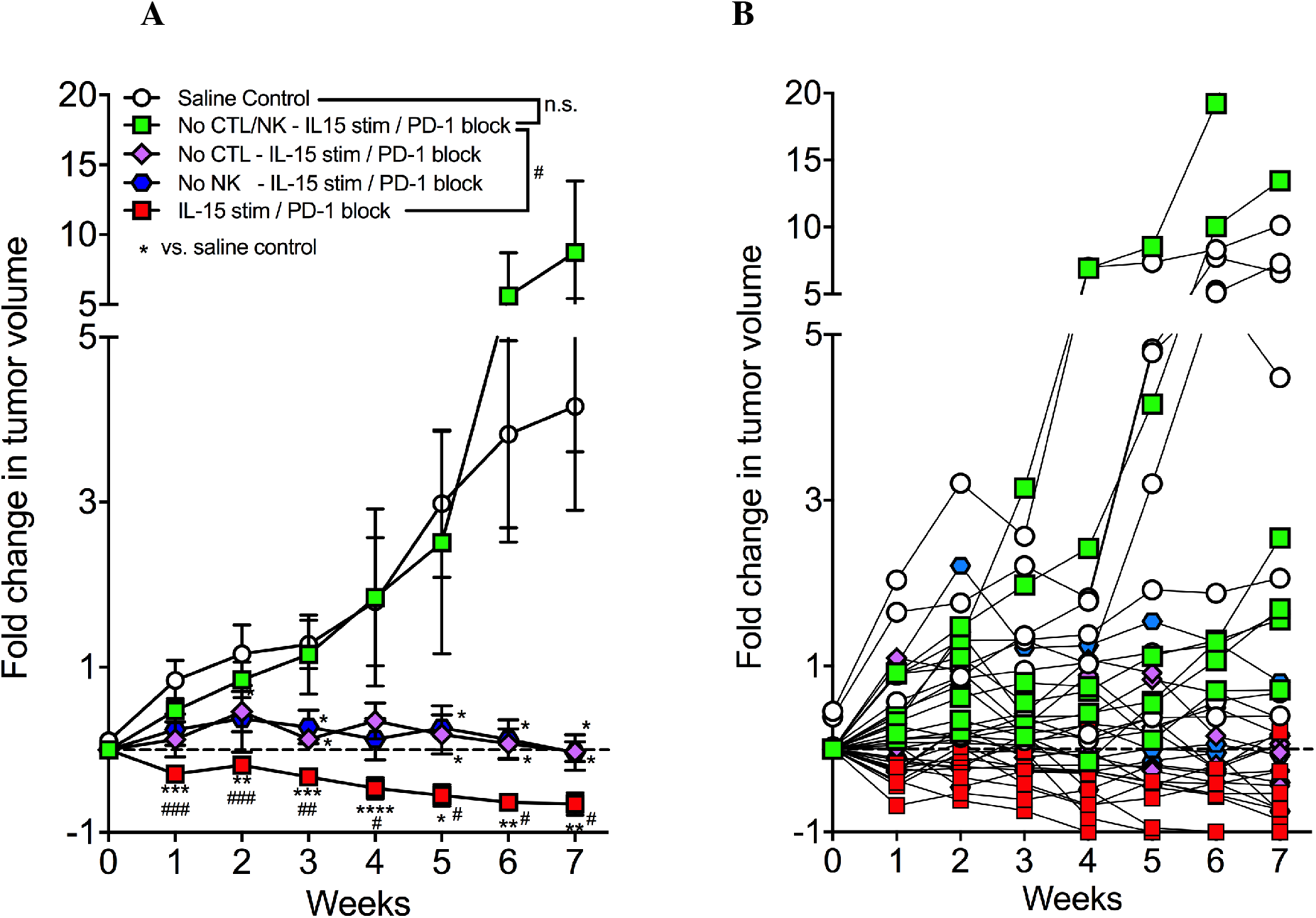
NK cells and CTLs are required for successful combination immunotherapy in LUAD-SIS-PDX model. (**A**) Fold change in tumor volume in LUAD-SIS-PDX mice treated with 100μg/mouse blocking antibody to PD-1 and 5μg/mouse IL-15 + 2.5μg/mouse IL-15RaFc by intra-peritoneal injection at day 0 and every seven days thereafter. Control mice received saline. Experiments were started when tumors were about 50 +/− 14 mm^3^ with no statistic size difference between experimental and control groups at day 0 and tumor volumes measured weekly. Three days prior to the start of PD-1 + IL-15/IL-15RaFC therapy (day -3) as well as on days 0, 7, 14, 21, 28, 35, 42, and 49. CTL and/or NK cells were depleted using 100μg/mouse of antibody specific to human CD8a (OKT-8) to deplete CTL, and/or antibody specific to NKp46 (9E2) to deplete NK cells, as indicated. Control mice received saline. (**A**) Mean fold change in tumor volume of each experimental or control group with standard error of the mean (**B**) Mean fold change in tumor volume of LUAD-SIS-PDX mouse with standard error of the mean. N = 6-9 LUAD-SIS-PDX mice per group from three genetically unrelated human donor cohorts. Data are represented as mean SEM; Multiple t-test, *,^#^p < 0.05; **,^##^ p < 0.01; ***,^###^ p < 0.001.

## Discussion

Current PDX models lack a human tumor-donor-matched, complete, functional and mouse-tolerant immune system (*53*–*57*). One PDX model relies on serial transplantation of solid tumors into NOD/SCID/IL2Rγ_c_-KO (NSG) mice to maintain genetic tumor diversity and allow for the evaluations of tumor genetics and chemotherapies (*66*). However an infusion of donor-matched or mismatched PBMC is required for immune system reconstitution, which results in the development of graft versus host disease (GvHD) within weeks of infusion (*53*). To circumvent GvHD, the second PDX model was generated by infusion of umbilical cord derived hematopoietic stem cells (UB-HSC) into immune compromised mice before tumor is engrafted, resulting in the development of mouse tolerant T cells (*53*). However, while GvHD was avoided, UB-HSC derived T cells were not restricted to human HLA due to the lack of a human thymus, resulting in artifactual CTL responses. Further, NK cells may or may not be inhibited by tumor-expressed HLA, depending on the nature of UB-HSC-NK cell expressed Killer cell Immunoglobulin-like Receptors (KIR), resulting in either artificial inhibition or augmentation of NK cell anti-tumor responses (*53*). These hurdles limit the usefulness of such PDX models for the development and efficacy testing of novel or improved immunotherapy approaches to treat solid tumor.

Here, we introduce a novel PDX model of human solid tumor with a TIL-derived syngeneic complete functional and mouse-tolerant immune system. In our model, long-term immune reconstitution does not require the infusion of PBMC, or stem cells, thus avoiding GvHD or induction of allogeneic immune cells. Further, SIS-PDX model is independent of cytokine treatment of tumor-engrafted NSG mice, or the in vitro pre-expansion of human TILs. Instead, we achieve long-term immune reconstitution by implantation of an undisrupted and treatment-naïve piece of LUAD and by allowing TIL to repopulate the periphery of the recipient NSG mouse over time. About 70 percent of transplanted LUAD-SIS-PDX and about 80 percent PDAC-SIS-PDX mice harbor tumors and reconstitute long-term with tumor fighting-effector and tumor-associated regulatory cells. In reconstituted LUAD- and PDAC-SIS-PDX mice, CTL and NK cells resemble the human tumor-donors’ exhausted effector cells in phenotype and persist for many months post engraftment. Consistently, CTL and NK cell numbers within engrafted tumors inversely correlate with tumor size, while the expression of PD-ligands on NK cells correlate with tumor growth. Persistence of syngeneic immune cells in LUAD-SIS-PDX mice allows immunotherapy and mechanistic studies, whereby we show that IL-15-stimulation significantly augments ICI preventing treatment resistance. Interestingly, CTL and NK cells are both targets of this combination immunotherapy and are sufficient and equally required for tumor eradication. As such, our novel pre-clinical SIS-PDX-model is a valuable resource for powerful mechanistic and therapeutic studies for the eradication of solid tumor.

We have made the discovery that the expression of PD family members is tissue- and cell specific: We found that most lung- and LUAD-resident NK cells express PD-1 as well as PD-L1 and/or PD-L2. However, the frequency of PD-1, PD-L1 and PD-L2-expressing NK cells is significantly higher in lung tumor compared to donor-matched non-malignant lung tissue. In contrast, the same donor’s PBMC-derived NK cells expressed neither PD-1 nor PDL-1 or PDL-2. Similarly, T cells expressed PD-1 and/or PD-L2, but not PD-L1 when resident in both control lung tissue and LUAD, and PD-1 and/or PD-L2 expressing T cells were more frequent in tumor tissue compared to lung-tissue. However, also similar to our data on NK cells, most PBMC-derived T cells did not express PD-1 or either ligand. We cannot formally exclude the possibility that immune cell phenotypes in the lung and pancreatic tissues were somehow affected by adjacent tumors, however, whether PD family members are expressed constitutively on immune cells in a tissue specific manner deserves further study. Should PD members be tissue specifically expressed on T and NK cells, then studies using PBMC as a sole control tissue may not be able to distinguish between tissue-resident and tumor-specific phenotypes.

A major advance of our novel PDX model is the long-term engraftment of a syngeneic TIL-derived complete functional and mouse-tolerant immune system, especially as homeostatic expansion did not cause any GvHD, nor did it rescue CTL and NK cells from tumor-exhaustion, and residual immune cell anti-tumor activity failed to prevent tumor growth. As such, our SIS-PDX model resembles a cancer patient’s tumor-exhausted immune system and enables the testing of high-risk immunotherapy approaches, as well as in depth mechanistic studies of anti-tumor immune responses. While several currently ongoing clinical trials are evaluating the efficacy of ICI monotherapies in localized and locally advanced solid tumor types, including NSCLC (NCT03425643; NCT02998528), ICI monotherapies are approved as a therapy for stage III NSCLC only after chemo-irradiation (*67*), and not as a primary treatment. In line with our long-term goal to use SIS-PDX mice to develop effective primary combination immunotherapies, we examined whether the addition of IL-15 stimulation to PD-1 blockade unleashes a strong anti-tumor response in NSCLC-SIS-PDX mice by therapeutically targeting tumor-exhausted CTL and NK cells in LUAD-SIS-PDX mice. To our surprise, IL-15 stimulation was far superior to PD-1 blockade, which temporarily blocked tumor growth before tumor quickly escaped from PD-1 blockade and grew at a rate similar to untreated tumor. In contrast, IL-15 stimulation resulted in tumor attach in all animals and reduction of tumor size. Importantly, the combination of IL-15 stimulation and PD-1 blockade elicited a strong anti-tumor response in all treated PDX mice, demonstrating that IL-15 stimulation prevents tumor escape to ICI and augments anti-tumor immunity. These data may be especially important for the treatment of solid tumors for which ICI is of limited benefit, such as neo-antigen-poor tumors such as PDAC. In PDAC, CTL exhaustion can be prevented or reversed by PD-1 and PD-L1 blockade, however, MHC-I-null pancreatic cancer cells emerge and escape CTL killing (*68*). This MHC I downregulation however may be exploitable for tumor attack, as NK cells, which, unlike CTL, sense the lack of MHC I expression and are potent tumor fighters responsive to IL-15, the therapy we tested here either by itself or in combination with PD-1 blockade. IL-15 stimulation to ICI is likely safe, as demonstrated by a recent Phase I clinical trial that established that IL-15 stimulation and PD-1 blockade is both safe for and beneficial to a portion of NSCLC patients who had failed conventional therapies (*49*). However, albeit understandably, this study (*49*) was conducted with a small number of patients for which IL-15 stimulation and PD-1 blockade was not their primary therapy, and no mechanistic studies could be performed.

To understand why our combination immunotherapy eradicates LUAD in SIS-PDX mice, we asked whether CTLs and/or NK cells were required for tumor eradication upon IL-15 stimulation and PD-1 blockade. Interestingly, we found that both cell types were equally required and their tumor-fighting actions synergistic to eradicate human solid tumor. Our findings demonstrate an unexpected plasticity of exhausted NK cells and CTLs in the tumor microenvironment that can be redirected therapeutically to jointly fight tumor in vivo, and agree with reports that identified NK cell- and CTL-cooperation as crucial for the control of solid tumors in mice (*8*, *9*), and a recent report on the effective augmentation of PD-L1-blockade-induced NK cell and CTL-mediated anti-tumor activities by IL-15 stimulation in murine models of solid tumor (*69*). In agreement with these data obtained in mice, our data demonstrates that IL-15 stimulation effectively activates human anti-tumor activity in NK cells and CTLs in combination with ICI. It is worth mentioning that immunotherapy elicited by NK cell contributions to tumor eradication may be even more significant than what our data suggests, as NK cell depletion with the commercially available antibody to NKp46 targets only NKp46-domain 1 expressing NK cells, leaving NKp46-domain 2-expressing NK cells intact. As such, future studies on the role of specific NK cell subsets in the rejection of solid tumor in response to immunotherapy are needed to understand the precise roles human NK cell subsets play in solid tumor eradication.

A significant benefit of our novel SIS-PDX model is that it allows the study of the human immune response to genetically diverse human tumors from both male and female human donors, in contrast to mouse tumor models, in which a specific mutation or combination of mutations resemble a subset of human tumors (*70*). It is therefore noteworthy our therapy generally worked well with any tumor donor and across individual PDX mice of a given cohort, given the expected genetic tumor heterogeneity within a single tumor and within the human cancer patient population, suggesting that this model may work well to study immunotherapy across a diverse human population. One limitation worth mentioning however is that orthotopic transplantation of solid tumor may not be possible for all tumor types. We transplanted both LUAD and PDAC into skin pockets of NSG mice but did not observe any metastases in our SIS-PDX model. However, PDAC can be transplanted orthotopically, and future studies will examine whether immune reconstitution is robust in orthotopically transplanted PDAC-SIS-PDX mice.

In summary, our new pre-clinical model provides a valuable resource for a multitude of mechanistic and therapeutic studies not previously possible due to the lack of an appropriate translational model. Using solid-tumor-SIS-PDX mice, we provide a strong pre-clinical evidence for the use of combination immunotherapy to treat solid tumor and highlight the importance of NK cell – CTL cooperation in therapies frequently thought to exclusively target T cells.

## Materials and Methods

### Study design

The goals of this study were to develop a novel PDX model in which (i) tumor and immune cells are from the same human donor, (ii) relevant human immune effector and suppressor cells are present in PDX mice and persist long-term, (iii) human immune cells resemble the patient’s tumor-exhausted immune system allowing tumor growth, (iv) the expression of checkpoint molecules and markers of exhaustion is preserved on tumor and immune cells, and (v) human immune cells do not cause GvHD. Having developed such a model, the next goal of this study was to develop a safe and effective immunotherapy preventing solid tumor immune escape from ICI and eliciting tumor eradication. Having developed such a therapy, the remaining goal of this study was to determine whether NK cells and / or CTL are required and sufficient for immunotherapy-elicited tumor eradication. The numbers of human donors and PDX mice per group per experiment and statistical analyses used to analyze data are indicated in the figure legends of each corresponding data figure.

#### Mice

Non-Obese-Diabetic / Severe Combined Immunodeficient / Interleukin 2 Receptor common gamma chain-deficient (NOD/SCID/Il2Rγc or NSG) mice were purchased from the Jackson Laboratory and bred at Baylor College of Medicine (BCM) under pathogen-free conditions in sterile microisolator cages. All mice were cared for in either the animal facilities of the Center for Comparative Medicine at Baylor College of Medicine (BCM), Houston, TX, or the animal facilities of The Scripps Research Institute, La Jolla, CA, and all protocols were approved by the Institutional Animal Care and Use Committees of the respective institutes and experiments performed in accordance to the Animal Welfare Act guidelines.

#### Research ethics approval for PBMC, adult LUAD, lung and PDAC tissue

All human peripheral blood and adult spleen samples were obtained by written informed consent and the protocol was approved for use by the National Institutes of Health (NIH), Baylor College of Medicines’ Institutional Review Board for the Protection of Human Subjects, and The Scripps Research Institutes’ Institutional Review Board and in accordance with the Declaration of Helsinki.

#### PBMC from LUAD and PDAC donors

To isolate PBMC, peripheral blood was diluted 1:1 with Ca^++^, Mg^++^ free phosphate buffered saline (PBS) and layered over Ficoll-Pacque (GE Healthcare), centrifuged at 2,000 rpm with no brake for 20 minutes at room temperature (RT) according to the manufacturer’s instructions. The lymphocyte layer was removed, washed, counted, and immune cells stained with the indicated antibodies and analyzed using multiparametric flow cytometry.

#### Tumor transplantation

Freshly dissected small-undisrupted pieces of LUAD or PDAC were sectioned into areas of 2-4 mm^3^, which were transplanted subcutaneously into six to eight-week NSG mice that were sex-matched to the human tumor donor. A digital caliper was used to measure tumor sizes weekly after transplant. Tumor volumes were calculated using the following formula: ½ (L × W^2^) where L is length, W is width of the tumor.

#### Processing of tissues

PDX mice were euthanized when their tumors exceeded 100 mm^3^ in volume, or at the experimental time points as indicated in the results section of this manuscript. Blood was collected from PDX mice either by submandibular bleed (live animals) or by cardiac puncture after euthanasia, and immune cells isolated using Ficoll-Plaque^™^ (GE Healthcare) gradient centrifugation. Tumors and spleens were excised from PDX mice and single cell suspensions generated by mechanical disruption of tissues followed by filtering of cell suspensions through 40μm nylon mesh. Immune cells were separated by density gradient centrifugation using Ficoll-Plaque^™^. Layers of immune cells were separated from gradients and washed with PBS / 2% FBS before proceeding with antibody staining.

#### Flow cytometry

Immune cells were treated with human- and mouse-specific Fc Block^™^ (BD Biosciences) at room temperature for 10 minutes to prevent non-specific antibody staining. Immune cells were then stained using indicated combinations of fluorescent antibodies listed in **Supplemental Table 1**. Extracellular staining was performed for 30 minutes at 4°C in the dark. For *FOXP3* staining, Transcription Factor Staining Buffer Kit (Tonbo Biosciences) was used to fix and permeabilize cells before treating cells with Fc Block and subsequent intracellular staining. As controls, Fluorescence Minus One (FMO) controls were generated using human and murine PBMCs to distinguish between positive and negative cell populations. Gating was performed on life single cells that were negative for murine CD45 and positive for human CD45. Gating for specific cell types is described in the results section of this manuscript. Samples were analyzed on either an LSR Fortessa (BD Biosciences) or a 5-laser Aurora (Cytek) and raw data analyzed further using FlowJo (version 10.4.2).

#### Histopathology staining

Tumors excised from PDX mice were fixed in 10% formalin, before Hematoxylin and eosin (H&E) staining for histopathological analysis was performed fee for service by the Comparative Pathology Laboratory, Center for Comparative Medicine, Baylor College of Medicine, Houston, TX, or by the Pathology Lab Services of The Scripps Research Institute, La Jolla, CA.

#### Immunotherapy of PDX mice

Six to eight-weeks post-transplantation, PDX mice with tumors about 50 mm^3^ in size were injected intraperitoneally (i.p.) once a week every week with IL-15 Receptor alpha Fc Chimera (R&D; 2.5μg/mouse) and recombinant human IL-15 (Biolegend, 5 μg/mouse). For immune blockade of PD-1, PDX mice were injected i.p. weekly with 100μg of anti-human PD-1 (Bioxcell, J116). All treatments were administered together on the same day.

#### NK cell and CTL depletion *in vivo*

In order to deplete human NKp46 domain 1 containing NK cells and CTL cells, PDX mice were injected i.p. with 100ug of antibodies against specific human NK cells expressing NKp46 Domain 1 (Biolegend, 9E2) and/or human CD8α (Bioxcell, OKT-8) three days prior to first treatment and concurrently with subsequent treatments.

#### Statistical analysis

All Statistical analyses were calculated using GraphPad Prism 7 (GraphPad Software). The unpaired Welch’s t-test was used to compare two groups of unpaired data. The paired t-test was used when comparing two groups of paired data. The Pearson correlation coefficient was computed to identify correlations. Data was presented along with mean ± SD or mean ± SEM as indicated. Statistical significance was represented as *p < 0.05; **p < 0.01; ***p < 0.001.

#### Study Approval

All human peripheral blood and adult LUAD, lung and PDAC tissues were obtained by written informed consent and the protocol was approved for use by the National Institutes of Health (NIH), Baylor College of Medicines’ Institutional Review Board (IRB) for the Protection of Human Subjects, as well as The Scripps Research Institutes’ Institutional Review Board and in accordance with the Declaration of Helsinki. For all animal experiments, all protocols were approved by the Institutional Animal Care and Use Committees (IACUC) of the respective institutes and experiments performed in accordance to the Animal Welfare Act guidelines.

## Author Contributions

Conceptualization – SP

Formal analysis – DTL, SAA, SP

Funding acquisition – SP, GvB, FK

Investigation – DTL, BB, GvB, SAA, MCZ, RN, SP

Methodology – DTL, BB, GvB, RN, SP

Project administration – SP

Resources – BB, GvB

Supervision – SP

Validation – DTL, SAA, RN, SP

Visualization – SP

Writing –SP, FK

## Acknowledgements

This project was done in collaboration with the Adrienne Helis Malvin Medical Research Foundation through its direct engagement in the continuous active conduct of medical research in conjunction with Baylor College of Medicine and the Natural Killer Cell Immunotherapy to Cure Lung Cancer Program (to SP and FK), the Department of Pathology and Immunology, the Department of Medicine and the Department of Surgery at Baylor College of Medicine. Additional support included The Scripps Research Institute’s unrestricted funds to SP. We thank Mr. Christopher Lanier, Mr. Hangqing Lin and Dr. Mayra Shanley for assistance with animal care and generation of PDX mice, and Ms. Martina Navarro Cagigas for her assistance with the coordination of PDAC samples.

## The following data and information can be found in the supplemental materials

**Supplemental Figure 1:** Long-term reconstitution of LUAD-SIS-PDX mice with human B cells and dendritic cells (DC).

**Supplemental Figure 2.** PD-L1 and PD-L2-expression by freshly resected and LUAD-SIS-PDX-derived tumor.

**Supplemental Figure 3.** Immune suppressive MDSC and M2 persistence in PDAC-SIS-PDX mice.

**Supplemental Figure 4.** PD-L1 and PD-L2-expression by freshly resected and SIS-PDX-derived PDAC.

**Supplemental Figure 5.** Human immune cells are required for successful PD-1 + IL-15/IL15Ra-Fc-chimera therapy.

---

**Supplemental Figure 1:**
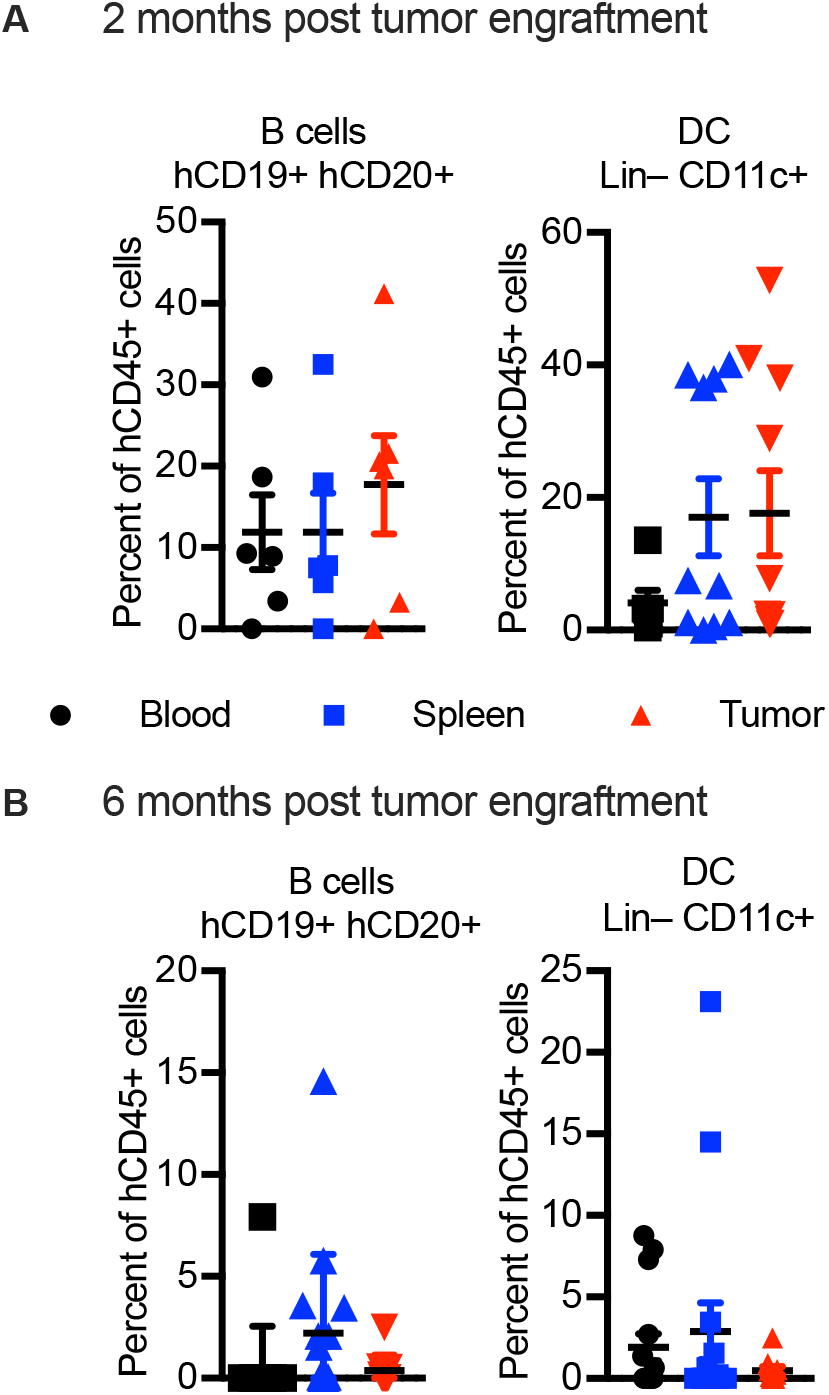
Long-term reconstitution of LUAD-SIS-PDX mice with human B cells and dendritic cells (DC). Frequencies of Human B cells (CD45+CD3−CD56−CD19+CD20+) and DC (CD45+CD3−CD56−CD19−CD11c+) in PBMC (“blood”), spleen and transplanted tumor of LUAD-SIS-PDX mice (**A**) 2- and (**B**) 6-months post tumor-engraftment, as determined by flow cytometry. Each dot represents one LUAD-SIS-PDX mouse, N= 10-15 animals per tissue.

**Supplemental Figure 2:**
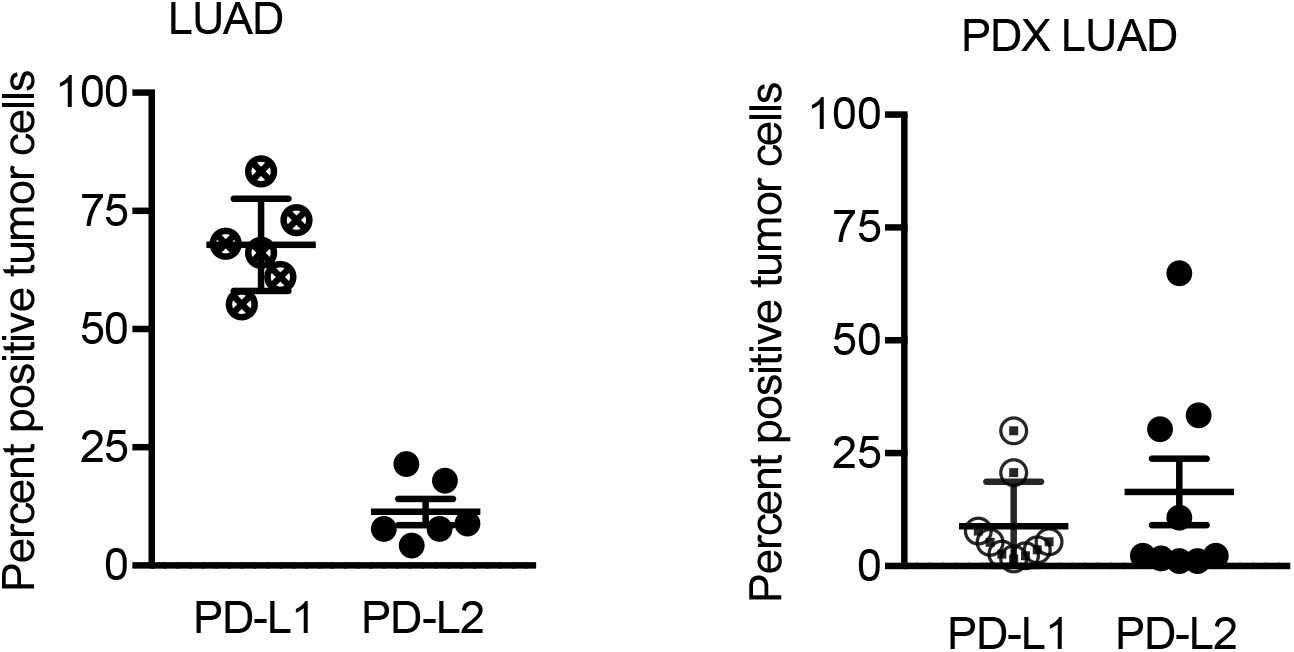
PD-L1 and PD-L2-expression by freshly resected and LUAD-SIS-PDX-derived tumor. Frequency of PD-L1 and PD-L2 expressing non-hematopoietic cells (CD45^−^) in freshly dissected or LUAD-SIS-PDX-derived tumor 4 months post-transplant as determined by flow cytometry. N = 6 freshly excised tumor tissues were obtained from genetically unrelated human donors and analyzed by flow cytometry, as well as N = 9 SIS-PDX mice from three genetically unrelated donor cohorts. Freshly excised and SIS-PDX derived tumor tissues were not from the same donor.

**Supplemental Figure 3:**
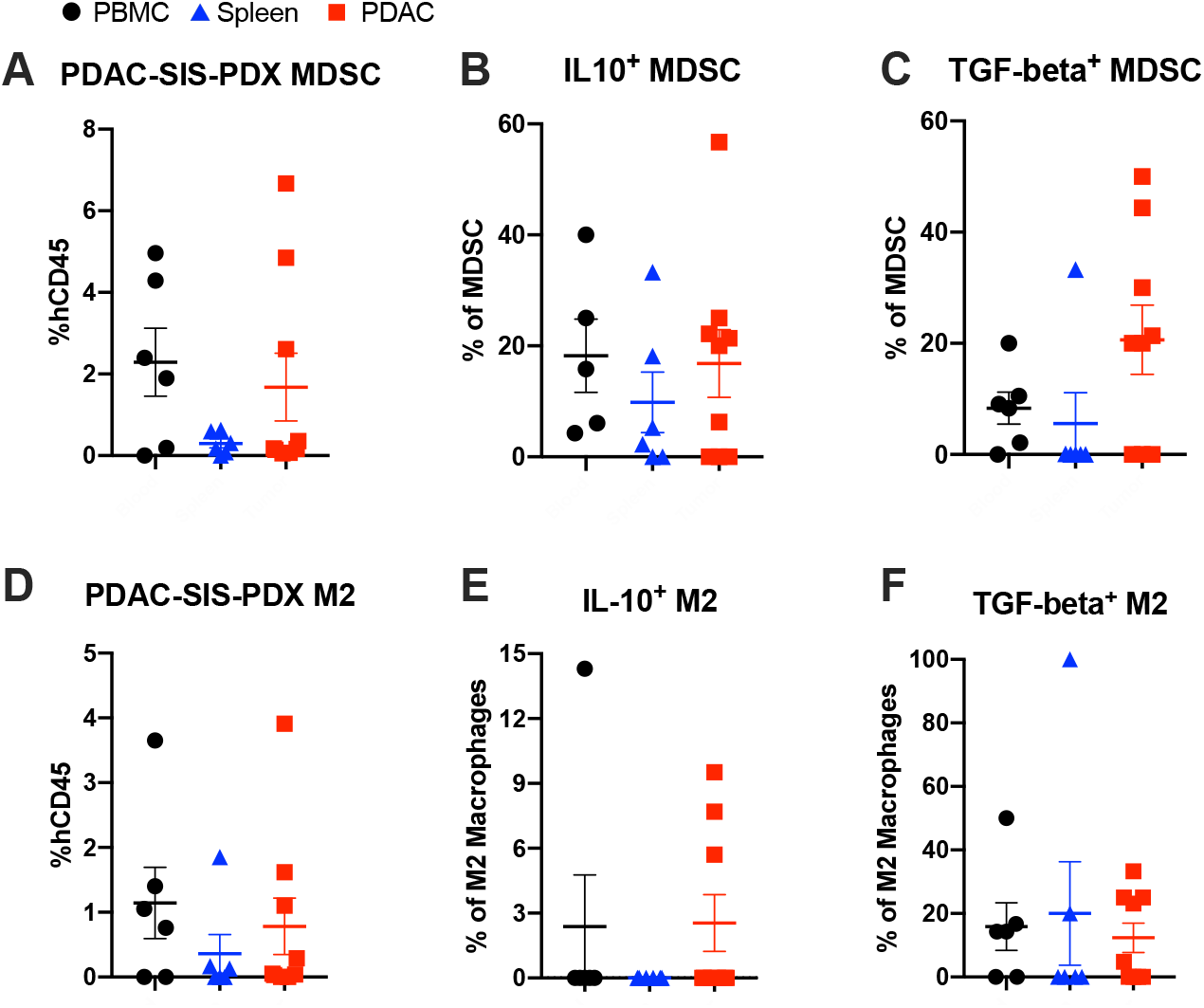
Immune suppressive MDSC and M2 persistence in PDAC-SIS-PDX mice. Frequency of (**A**) MDSC (human CD45^+^CD3^−^CD56^−^CD16^−^HLADR^low/−^ CD11b^+^CD33^+^), (**B**) IL-10 producing MDSC and (**C**) TGF-beta-producing MDSC, as well as (**D**) PDAC-SIS-PDX M2 macrophages (CD45^+^CD3^−^CD56^−^CD14^+^CD163^+^), (**E**) IL-10 producing M2 macrophages and (**F**) TGF-beta-producing M2 monocytes in PBMC, spleens and PDAC of PDAC-SIS-PDX mice 4-months post-transplant as determined by flow cytometry. N = 7 SIS-PDX mice from three genetically unrelated donor cohorts were analyzed.

**Supplemental Figure 4:**
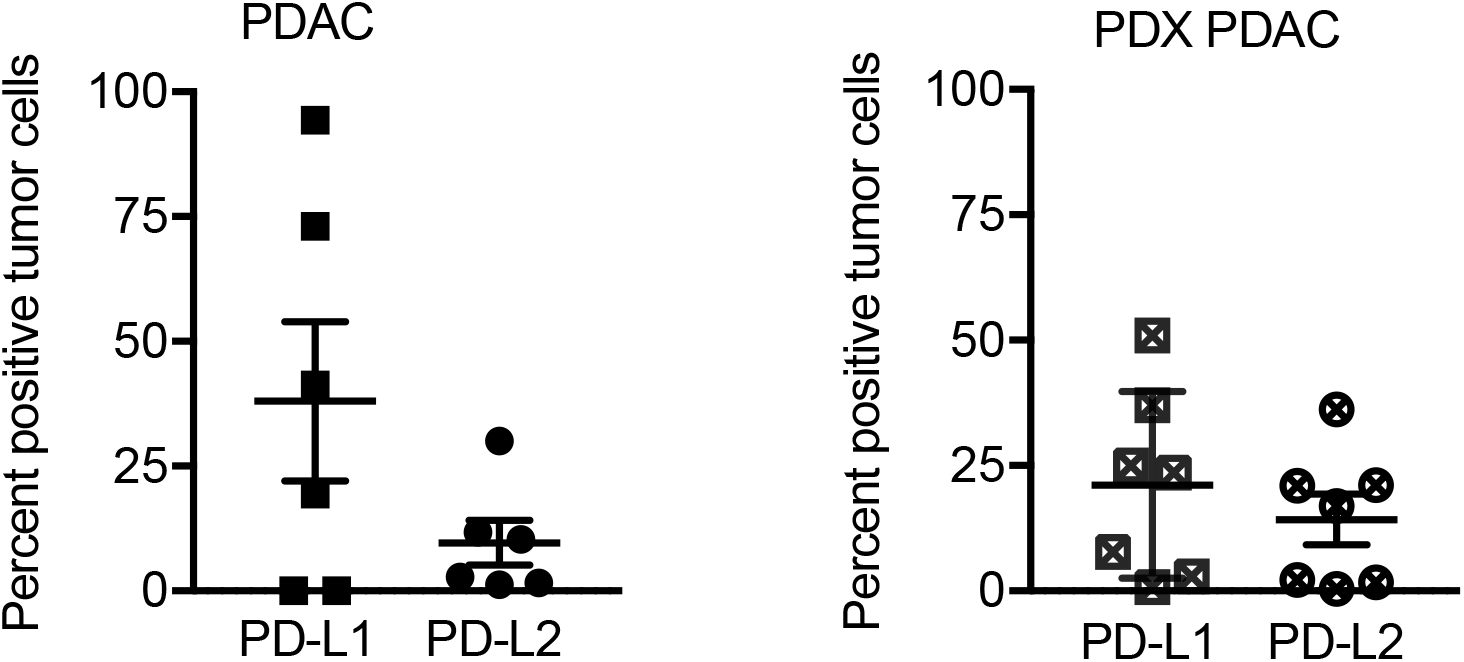
PD-L1 and PD-L2-expression by freshly resected and SIS-PDX-derived PDAC. Frequency of PD-L1 and PD-L2 expressing non-hematopoietic cells (murine CD45^−^ and human CD45^−^) in freshly dissected or PDAC-SIS-PDX tumor tissue 4 months post-transplant as determined by flow cytometry. N = 6 freshly excised PDAC tumor tissues were obtained from genetically unrelated human donors, and N = 7 PDAC-SIS-PDX mice from three genetically unrelated donor cohorts and analyzed by flow cytometry. Freshly excised and SIS-PDX derived tumor tissues were not from the same donor.

**Supplemental Figure 5:**
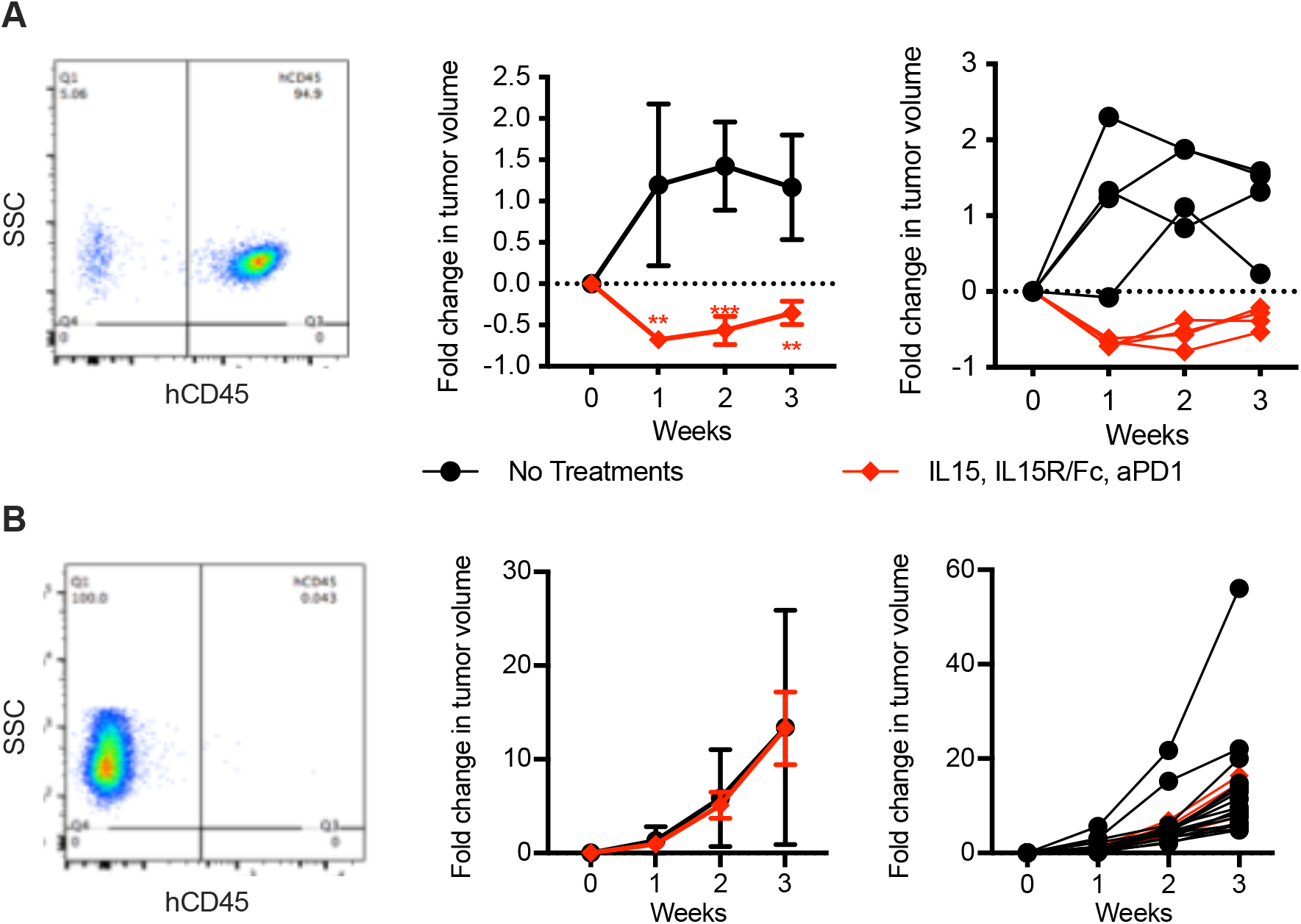
Human immune cells are required for successful PD-1 + IL-15/ IL15RaFc-chimera therapy. (**A**) Freshy harvested LUAD tumor tissue was transplanted into NSG mice, (**B**) while the other half was frozen at −80°C for two weeks before being transplanted into NSG mice. Flow cytometry of LUAD-SIS-PDX mice demonstrating (**A**) the presence of human immune cells (human CD45^+^) in recipients of fresh tumor tissue, compared to (**B**) the absence of human immune cells in recipients of frozen tumor tissue. LUAD-SIS-PDX mice were treated with either human IL-15 + human IL-15Ra Fc chimera + PD-1 blocking antibody or left untreated, and tumor volumes were recorded weekly. N = 4 – 16 LUAD-PDX mice per group from two genetically unrelated donor cohorts, thus N= 8-32 mice total per group). Data are represented as mean SEM; Multiple t-test, *p < 0.05; **p < 0.01; ***p < 0.001.

**Supplemental Table 1.**

| **Antibodies** | **Fluorophore** | **Clone** | **Source** | **Identifier** |
| --- | --- | --- | --- | --- |
| **Anti-human CD45** | Brilliant Violet 605™ | HI30 | Biolegend | 304042 |
| **Anti-human CD3** | Brilliant Violet 711™ | OKT3 | Biolegend | 317328 |
| **Anti-human CD3** | APC/Fire 750 | UCHT1 | Biolegend | 300470 |
| **Anti-human CD56** | Brilliant Violet 421™ | HCD56 | Biolegend | 318328 |
| **Anti-human CD56** | Brilliant Violet 570™ | HCD56 | Biolegend | 318330 |
| **Anti-human CD4** | Brilliant Violet 650™ | OKT4 | Biolegend | 317436 |
| **Anti-human CD4** | Brilliant Violet 750™ | SK3 | Biolegend | 344644 |
| **Anti-human CD8a** | Brilliant Violet 785™ | RPA-T8 | Biolegend | 301046 |
| **Anti-human CD11c** | PE/Cy7 | 3.9 | Biolegend | 301608 |
| **Anti-human CD11b** | PE/Dazzle™ | ICRF44 | Biolegend | 301348 |
| **Anti-human CD16** | Brilliant Violet 510™ | 3G8 | Biolegend | 302048 |
| **Anti-human CD16** | PE/Cy7 | 3G8 | Biolegend | 302010 |
| **Anti-human CD19** | Alexa Fluor® 700 | HIB19 | Biolegend | 302226 |
| **Anti-human CD20** | APC/Cy7 | 2H7 | Biolegend | 302314 |
| **Anti-human CD33** | FITC | HIM3-4 | Biolegend | 303304 |
| **Anti-human CD94** | PerCP Cy5.5 | DX22 | Biolegend | 305514 |
| **Anti-human CD163** | Brilliant Violet 711™ | GHI/61 | Biolegend | 333630 |
| **Anti-human PD-1** | Brilliant Violet 785™ | EH12.2H7 | Biolegend | 329930 |
| **Anti-human PD-1** | Brilliant Violet 480™ | EH12.1 | BD Biosciences | 566112 |
| **Anti-human PD-L2** | Alexa Fluor® 488 | 176611 | R & D | FAB1224G |
| **Anti-human PD-L2** | APC | MIH18 | Biolegend | 345508 |
| **Anti-human PD-L1** | Alexa Fluor® 700 | 130021 | R & D | FAB1561N |
| **Anti-human NKG2D** | Brilliant Violet 510™ | 1D11 | Biolegend | 320816 |
| **Anti-human NKG2D** | Super Bright 436 | 1D11 | ThermoFisher | 62-5878-42 |
| **Anti-human FOXP3** | Alexa Fluor® 647 | 150D | Biolegend | 320014 |
| **Anti-human FOXP3** | Alexa Fluor® 532 | PCH101 | ThermoFisher | 58-4776-42 |
| **Anti-human HLA-DR** | Brilliant Violet 650™ | L243 | Biolegend | 307650 |
| **Anti-human IL-10** | Alexa Fluor® 647 | JES3-9D7 | Biolegend | 501412 |
| **Anti-human TGF-beta** | PerCP/eFluor 710 | FN LAP | Thermofisher | 46-9829-42 |
| **Anti-mouse CD45** | PE/Cy5 | 30-F11 | Biolegend | 103110 |
| **Anti-mouse CD45** | PE | 30-F11 | Biolegend | 103106 |
| **Anti-mouse CD45** | PerCP/Cy5.5 | 30-F11 | Biolegend | 103132 |
| **Human Fc Block™** | N/A | N/A | BD Biosciences | 564220 |
| **Mouse Fc Block™** | N/A | 2.4G2 | BD Biosciences | 553142 |

